# Bioinformatic analysis of endogenous and exogenous small RNAs on lipoproteins

**DOI:** 10.1101/246900

**Authors:** Ryan M. Allen, Shilin Zhao, Marisol A. Ramirez Solano, Danielle L. Michell, Yuhuan Wang, Yu Shyr, Praveen Sethupathy, MacRae F. Linton, Gregory A. Graf, Quanhu Sheng, Kasey C. Vickers

**Author notes:** Co-first authors. Co-corresponding authors CORRESPONDING AUTHOR: Kasey C. Vickers, PhD, 2220 Pierce Ave. 312 Preston Research Building, Nashville, TN 37232. **ABBREVIATIONS** exRNA: extracellular RNAs HDL: high-density lipoproteins HMB: human microbiome project lncRNA: long non-coding RNA lncDR: lncRNA-derived sRNA LDL: low-density lipoproteins miscRNA: miscellaneous sRNA ncRNA: non-coding RNA NIH: National Institutes of Health nts: nucleotides osRNA: other sRNA PM: perfect match rDR: rRNA-derived sRNA, RPM, Reads Per Million total reads rRNA: ribosomal RNA sRNA: small RNAs snDR: snRNA-derived sRNA snoDR: snoRNA-derived sRNA snoRNA: small nucleolar RNA snRNA: small nuclear RNA SR-BI: scavenger receptor BI sRNA-seq: small RNA sequencing, tDR, tRNA-derived sRNA tRNA: transfer RNA yDR: Y RNA-derived sRNA 3’ UTR: 3’ untranslated regions.

## Abstract

To comprehensively study extracellular small RNAs (sRNA) by sequencing (sRNA-seq), we developed a novel pipeline to overcome current limitations in analysis entitled, “*Tools for Integrative Genome analysis of Extracellular sRNAs (TIGER)”.* To demonstrate the power of this tool, sRNA-seq was performed on mouse lipoproteins, bile, urine, and liver samples. A key advance for the TIGER pipeline is the ability to analyze both host and non-host sRNAs at genomic, parent RNA, and individual fragment levels. TIGER was able to identify approximately 60% of sRNAs on lipoproteins, and >85% of sRNAs in liver, bile, and urine, a significant advance compared to existing software. Results suggest that the majority of sRNAs on lipoproteins are non-host sRNAs derived from bacterial sources in the microbiome and environment, specifically rRNA-derived sRNAs from Proteobacteria. Collectively, TIGER facilitated novel discoveries of lipoprotein and biofluid sRNAs and has tremendous applicability for the field of extracellular RNA.

## Introduction

High-throughput small RNA sequencing (sRNA-seq) is a state-of-the-art method for profiling sRNAs, and is widely-used across many disciplines. Although many software are currently available for sRNA-seq data analysis, most fail to meet the present demands for the study of host and non-host sRNAs across diverse RNA classes. This is particularly important for the investigation of extracellular RNA (exRNA), which have recently been found to be heterogeneous pools of host (e.g. human) and non-host (e.g. bacteria) sRNAs^1–3^. Furthermore, individual sRNA classes harbor distinct features, e.g. miRNA 3’ non-templated additions (NTA)^4–6^, and these features each require unique strategies for alignments and quantification. A key objective for data analysis is to account for all reads in the sRNA-seq dataset, and current approaches to sRNA profiling now require sophisticated analysis strategies. Therefore, we developed a novel data analysis pipeline entitled, “***T****ools for* ***I****ntegrative* ***G****enome analysis of* ***E****xtracellular s****R****NAs (****TIGER***).” This pipeline integrated host and non-host sRNA analysis through both genome and database alignments, and greatly improved the ability to account for a larger number of reads in sRNA-seq datasets. The TIGER pipeline was designed for the study of lipoprotein sRNAs; however, it has great applicability to all sRNA-seq studies.

The most extensively studied class of sRNAs is microRNAs (miRNA)^7^ and many sRNA-seq analysis tools are limited to only miRNA quantification^8^. In addition to miRNAs, many other classes of sRNAs are present in sRNA-seq datasets^9^. These include sRNAs derived from parent transfer RNAs (tRNA), ribosomal RNAs (rRNA), small nucleolar RNAs (snoRNA), small nuclear RNAs (snRNA), long non-coding RNAs (lncRNA), Y RNAs, and several other miscellaneous non-coding RNAs^10, 11^. For consistency in nomenclature, here, we will refer to these novel sRNA classes as tRNA-derived sRNAs (tDR), rRNA-derived sRNAs (rDR), lncRNA-derived sRNAs (lncDR), snRNA-derived sRNAs (snDR), snoRNA-derived SRNAs (snoDR), Y RNA-derived sRNAs (yDR) and other miscellaneous sRNAs (miscRNA). Outside of miRNAs and tDRs, the biological function(s) of these other endogenous sRNAs are unknown^11, 12^. Nevertheless, similar to miRNAs, many of these endogenous sRNAs are present in biological fluids and hold great potential as disease biomarkers or intercellular communication signals; however, tools for their analysis in sRNA-seq datasets are very limited^13, 14^.

In plasma and other biofluids, exRNAs are carried by extracellular vesicles (EV), lipoproteins, and ribonucleoproteins, which protect exRNAs against RNase-mediated degradation^15, 16^. Previously, we reported that lipoproteins - low-density lipoproteins (LDL) and high-density lipoproteins (HDL) - transport miRNAs in plasma, and lipoprotein miRNA signatures are distinct from exosomes^17^. Using real-time PCR-based TaqMan arrays, we further identified HDL-miRNAs that were significantly altered in hypercholesterolemia and atherosclerosis^17^. Currently, it is unknown if lipoproteins transport other sRNAs in addition to miRNAs. In a previous study, we reported that HDL transfer miRNAs to recipient cells and this process is regulated by HDL’s receptor, scavenger receptor BI (SR-BI), in hepatocytes^17^. SR-BI is a bidirectional transporter of cholesterol and a critical factor in reverse cholesterol transport pathway in which HDL returns excess cholesterol to the liver for excretion to bile. Currently, it is unknown if miRNAs, and potentially other sRNAs, on lipoproteins follow cholesterol and are transported to the liver for secretion to bile. Likewise, as SR-BI can export cholesterol to HDL, it is unclear whether SR-BI directly influences sRNAs on lipoproteins or in biofluids.

To demonstrate the power of the TIGER pipeline, we present results from our comprehensive analysis of lipoprotein sRNAs by high-throughput sRNA-seq. Using TIGER, we found that lipoproteins transport a wide-variety of host and non-host sRNAs, most notably, HDL and APOB particles transport non-host bacterial tDRs and rDRs. TIGER analysis was also used to demonstrate that lipoprotein sRNA signatures were distinct from liver, bile, and urine for host sRNAs. Moreover, TIGER analysis was used to determine the role of SR-BI in the regulation of exRNAs on lipoproteins and in biofluids. At the parent RNA level, SR-BI-deficiency had minimal impact on sRNA levels; however, by organizing sRNAs at the individual fragment level, we found that loss of SR-BI in mice resulted in significant changes to specific sRNA classes in different sample types. TIGER was designed to overcome many of the barriers and challenges in sRNA-seq analysis, particularly for exRNA, and its application uncovered many novel observations for sRNAs on lipoproteins and in liver and biological fluids.

## Results

### Lipoproteins transport distinct miRNA signatures

The full-compendium of exRNAs on lipoproteins has not been investigated and an unbiased approach to identifying and quantifying sRNAs on lipoproteins was warranted. To address this gap, high-throughput sRNA-seq was used to profile sRNAs on HDL and apoB-containing particles (APOB) purified from mouse plasma by size-exclusion chromatography (**Figure 1**-**Figure Supplement 1A-C**), and lipoprotein profiles were compared to mouse liver, bile, and urine. For host sRNAs, the TIGER pipeline prioritized annotated sRNAs in ranking order; miRNAs, tDRs, rDRs, snDRs, snoDRs, yDRs, lncDRs, miscellaneous sRNAs (miscRNA), and unannotated host genome sRNAs (**Figure 1**). sRNAs can be normalized by reads per million total reads (RPM) or reads per million class reads, e.g. total miRNA reads (RPM miR). To compare miRNA content between groups, real-time PCR was performed for 9 miRNAs across all samples and correlations between PCR results and sRNA-seq results based on each normalization method were compared by rank correlations. For these data, normalization by RPM (R^2^=0.45) showed a higher correlation between PCR and sequencing results than RPM miR (R^2^=0.17) (**Figure 2A**, **Figure 2** – **Source Data 1**) Lipoproteins, specifically APOB particles, were found to have less miRNA content, as reported by total miRNA counts (RPM), than livers which had the largest fraction of miRNAs per total reads (RPM) (**Figure 2B**). To compare miRNA signatures across sample types, Principal Coordinate Analysis (PCoA) was used, and lipoprotein and biofluid clusters were distinct from livers (**Figure 2C**). To quantify differences in the homogeneity of the miRNA profile multivariable distributions (miRNAs) within each groups, PERMANOVA tests were performed, and miRNA profiles of lipoproteins (HDL and APOB) and biofluids (bile and urine) were significantly different than livers (wild-type, WT, mice) – APOB (F=9.57, p=0.001), HDL (F=7.11, p=0.001), bile (F=5.56, p=0.001), and urine (F=8.42, p=0.001) (**Figure 2** – **Source Data 2**). Next, beta-dispersion tests were used to determine that lipoprotein (high-dispersions) and biofluid (high-disperisons) samples were significantly (ANOVA P<0.05) more dispersed (less consistent) than livers (low-dispersion) – APOB (F=31.03, p<0.0001), HDL (F=23.20, p<0.0001), bile (F=17.09, p<0.0001), and urine (F=15.47, p<0.0001) (**Figure 2C**). To further compare miRNA signatures between groups, high-end analyses were performed using hierarchical clustering and correlations of group means. Lipoprotein profiles clustered separately from liver and biofluids, and lipoproteins displayed high correlations between HDL and APOB groups and modest correlations to liver, bile, and urine groups (**Figure 2D**). These results suggest that HDL and APOB transport unique miRNA signatures that are distinct from liver with decreased homogeneity and increased dispersion.

**Figure 1.**
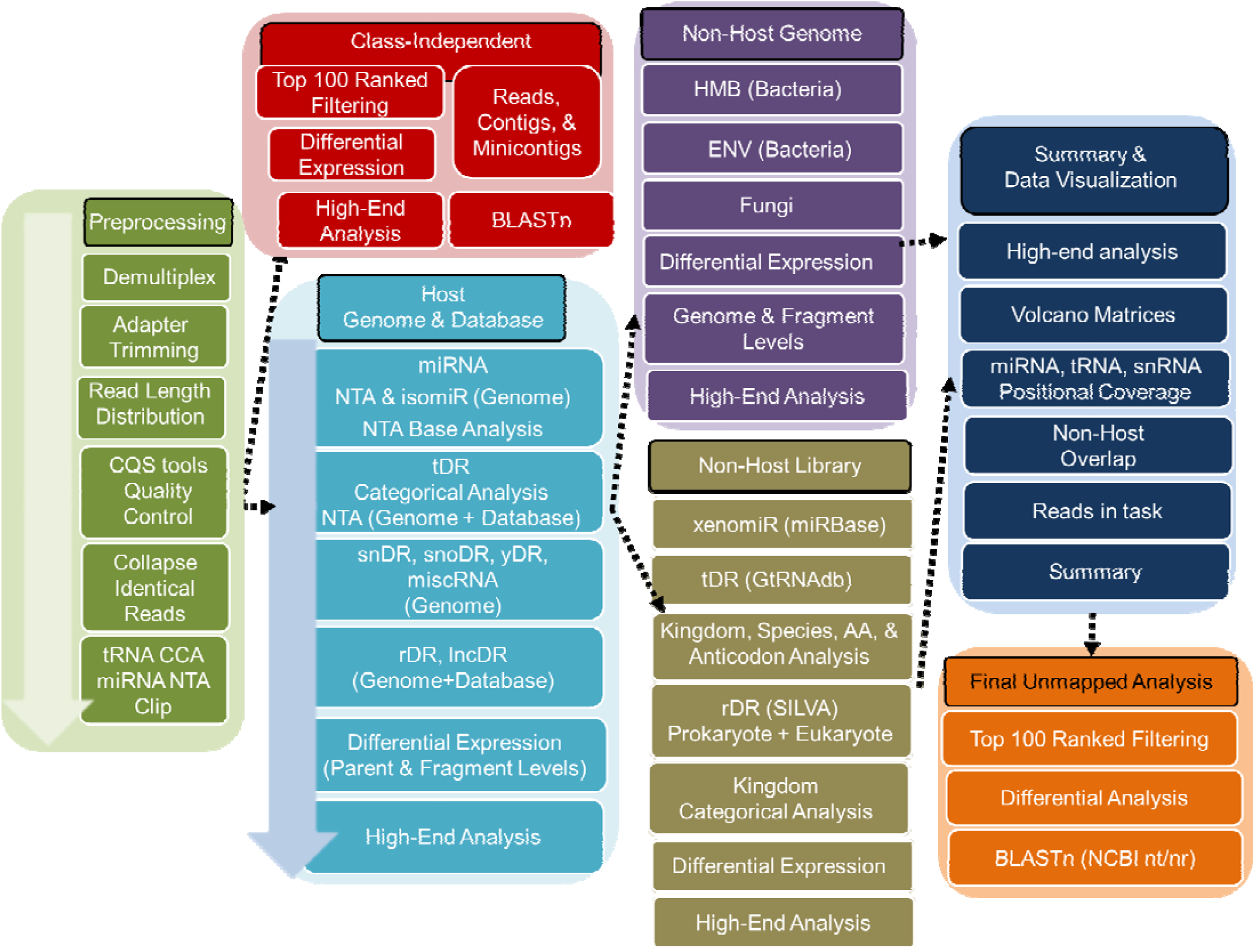
Schematic of the TIGER sRNA-seq analysis workflow.

**Figure 2.**
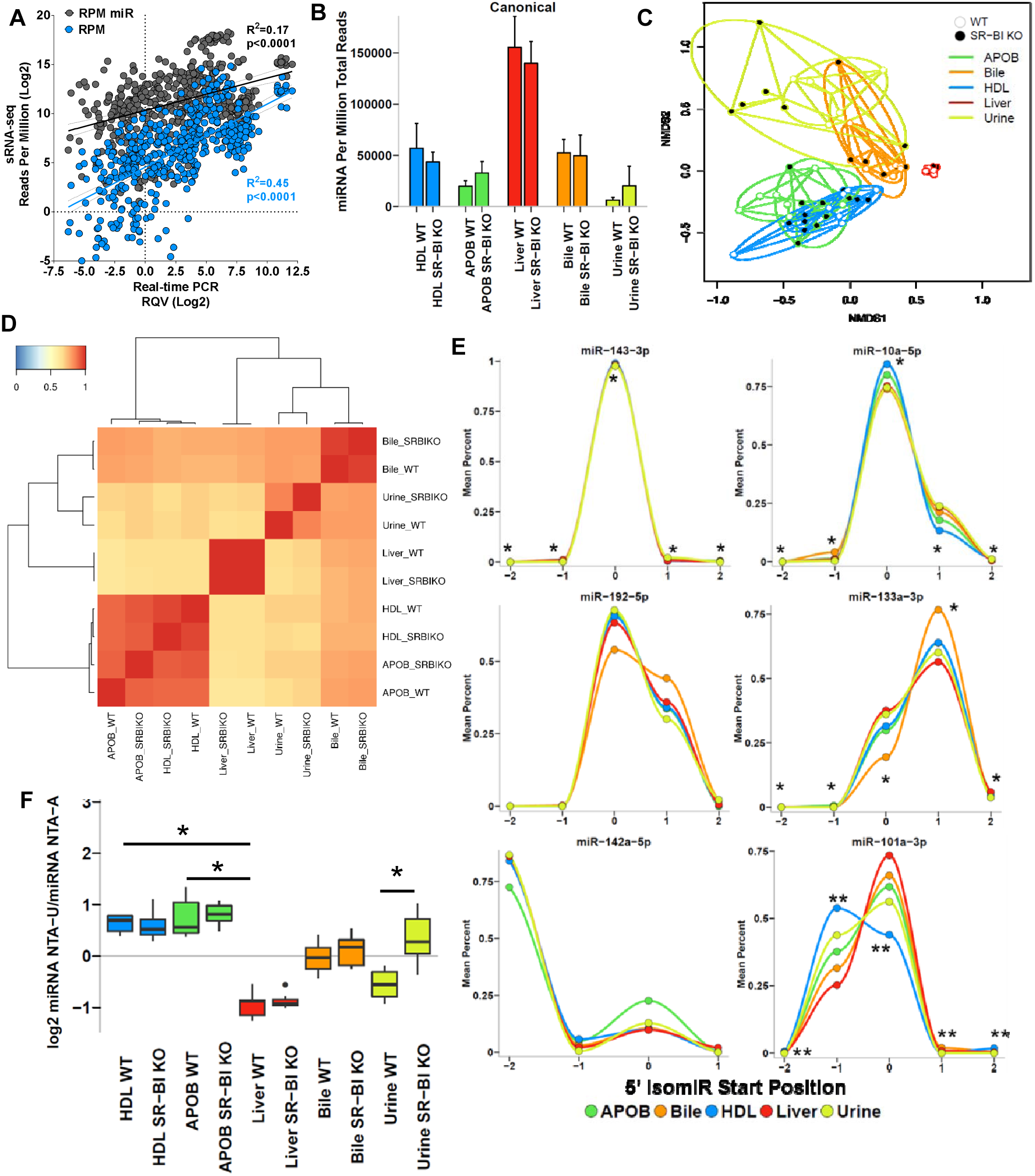
Host miRNAs on lipoproteins have distinct features compared to liver. WT, wild-type mice; SR-BI KO, Scavenger receptor BI Knockout mice (*Scarb1*^*-/-*^). (**A**) Correlation of sRNA-seq reads per million total reads (RPM, blue) and miRNA reads (RPM miR, gray) to real-time PCR relative quantitative values (RQV). Spearman correlation. HDL, APOB, liver, bile, and urine samples, N=66. (**B-F**) Results from sRNA-seq analysis. HDL WT, N=7; HDL SR-BI KO N=7; APOB WT, N=7, APOB SR-BI KO N=7; Liver WT, N=7; Liver SR-BI KO, N=7; Bile WT, N=7; Bile SR-BI KO, N=6; Urine WT, N=5; Urine SR-BI KO, N=6. (**B**) Abundance of canonical miRNAs. Mean ±S.E.M. (**C**) Principal Coordinate Analysis (PCoA) of canonical miRNA profiles for samples from WT (empty circles) and SR-BI KO (filled circles) mice. NMDS1, Non-metric multidimensional scaling. (**D**) Heatmap of hierarchical clustered pairwise correlation (Spearman, R) coefficients between group means for canonical miRNAs. (**E**) Start position analysis of 5’ miRNA variants (isomiR) for combined (WT and SR-BI KO) mouse samples. (**F**) Ratio of non-templated U (poly-uridylation) to A (poly-adenylation) for miRNAs. Mean ±S.E.M. One-way ANOVA. *p<0.05; **p<0.01

Due to imprecise cleavage of miRNAs from precursor miRNA hairpins^18–20^, one miRNA locus can produce multiple isoforms, termed isomiRs, which can differ by one or two nts at the 5’ start position. Consequently, the canonical miRNA “seed” sequence is altered, potentially conferring recognition of different mRNA targets^19–21^. Therefore, it is important that miRNA analysis includes quantification of isomiRs and all samples in our study contained 5’ isomiRs, the largest fraction was found on HDL (8.42%) followed by urine (7.2%), APOB (6.53%), bile (4.54%), and liver (4.34%) (**Figure 2**-**Figure Supplement 1A,B**). In addition, we found specific examples of miRNAs with different 5’ terminal start positions than their reported canonical forms, e.g. miR-142-5p (-2), miR-133a-3p (+1) and miR-192-5p (+1), and these patterns were consistent across all sample types (**Figure 2E**). Most interestingly, we found evidence that miRNAs may be partitioned to cellular and extracellular pools by their isomiR forms, as lipoproteins and biofluids contained significantly more 5’ (-1) isomiRs of miR-101a-3p than liver samples (**Figure 2E**). Mature miRNAs also harbor extensive variability on their 3’ terminal ends due to imprecise processing and NTAs, e.g. extra non-genomic 3’ nts added by cytoplasmic nucleotidyltransferases^6, 22^. A substantial fraction of miRNAs (17-32%) across all sample types were modified with NTAs (**Figure 2**–**Figure Supplement 1C**). As a percentage of total miRNAs, APOB particles contained significantly more miRNAs harboring non-templated additions (NTA) than liver samples (**Figure 2**-**Figure Supplement 1B,C**). A previous study proposed that poly-uridylation (NTA-U) was increased on extracellular miRNAs (released in exosomes) and miRNA poly-adenylation (NTA-A) was associated with cellular retention^23^. To determine if lipoproteins and/or biofluids are similarly enriched for poly-uridylation, NTA patterns were compared between groups, and similar to exosomes, HDL and APOB samples were indeed observed to be significantly enriched with NTA-U compared to liver samples which were enriched with NTA-A (**Figure 2F**). Intriguingly, extracellular miRNAs in bile and urine from WT mice were not enriched for either NTA; however, in urine, loss of SR-BI (KO mice) was found to significantly increase the NTA-U/A ratio (**Figure 2F**). Collectively, these results demonstrate that miRNAs on lipoproteins are distinct for many features from hepatic miRNAs, including 5’ isomiRs and 3’ NTAs.

### Lipoproteins transport many classes of host sRNAs

Most, if not all, non-coding RNAs are processed to smaller fragments creating an enormously diverse pool of sRNAs in cells and extracellular fluids^12^. To determine if lipoproteins also transport non-miRNA sRNAs and to compare annotated host sRNAs across sample types, reads were aligned to the host (mouse) genome, as well as to mature transcripts for specific RNA classes with genes containing introns, e.g tRNAs and rRNAs^24^. For liver samples, the most abundant class of sRNAs was rDRs, which were predominantly 42-45 nts in length (**Figures 3A,B**). rDRs were also present on HDL and APOB particles; however, their lengths were variable (**Figures 3A,C,D**). We also detected snoDRs (57-64 nts in length) in livers; however, snoDRs were largely absent from lipoproteins and biofluids, suggesting that the liver and other tissues may not export this class of sRNAs to lipoproteins or into bile or urine (**Figures 3A-F**). Both lipoproteins and biofluids contained tDRs 28-36 nts in length which suggests that these sRNA are likely tRNA-derived halves (tRHs), a sub-class of tDRs approximately 31-35 nts in length (**Figures 3A,C,D**)^42, 47^. Most tDRs on lipoproteins and in biofluids aligned to the 5’ halves of parent tRNAs, particularly amino acid anticodons for glutamate (GluCTC), glycine (GlyGCC), aspartate (AspGTC), and valine (ValCAC) (**Figure 4A**, **Figure 4**-**Figure Supplement 1**). Strikingly, 68.9% of tDR reads on HDL and APOB particles from WT mice aligned to the parent tRNA GluCTC (**Figure 4A**, **Figure 4**-**Figure Supplements 2A,B**). A key feature of the TIGER pipeline is the ability to analyze sRNAs based on their parent RNAs or individually as fragments. At the parent level for tDR signatures, all groups overlapped; however, at the fragment level, tDR signatures for lipoproteins, bile, and urine were found to be clearly delineated from livers (**Figures 4B,C**). These results were supported by PERMANOVA analysis which indicated that lipoprotein and biofluids were significantly distinct from liver based on tDR fragments: APOB (F=5.32, p=0.001), HDL (F=2.94, p=0.014), bile (F=10.22, p=0.001), and urine (F=7.08, p=0.001) (**Figure 2**-**Source Data 2**). Hierarchical clustering and correlation analyses further support that the profile of individual fragments, rather than parent tRNAs, define tDR expression across sample types (**Figure 4**-**Figure Supplements 3A,B**). Strikingly, this pattern where groups are defined by organizing sRNAs based on individual fragments instead of parent RNAs was consistent for other host sRNAs, including rDRs and snDRs (**Figure 4**-**Figure Supplements 3C-F,4A-F**; **Figure 2**-**Source Data 2**).

**Figure 3.**
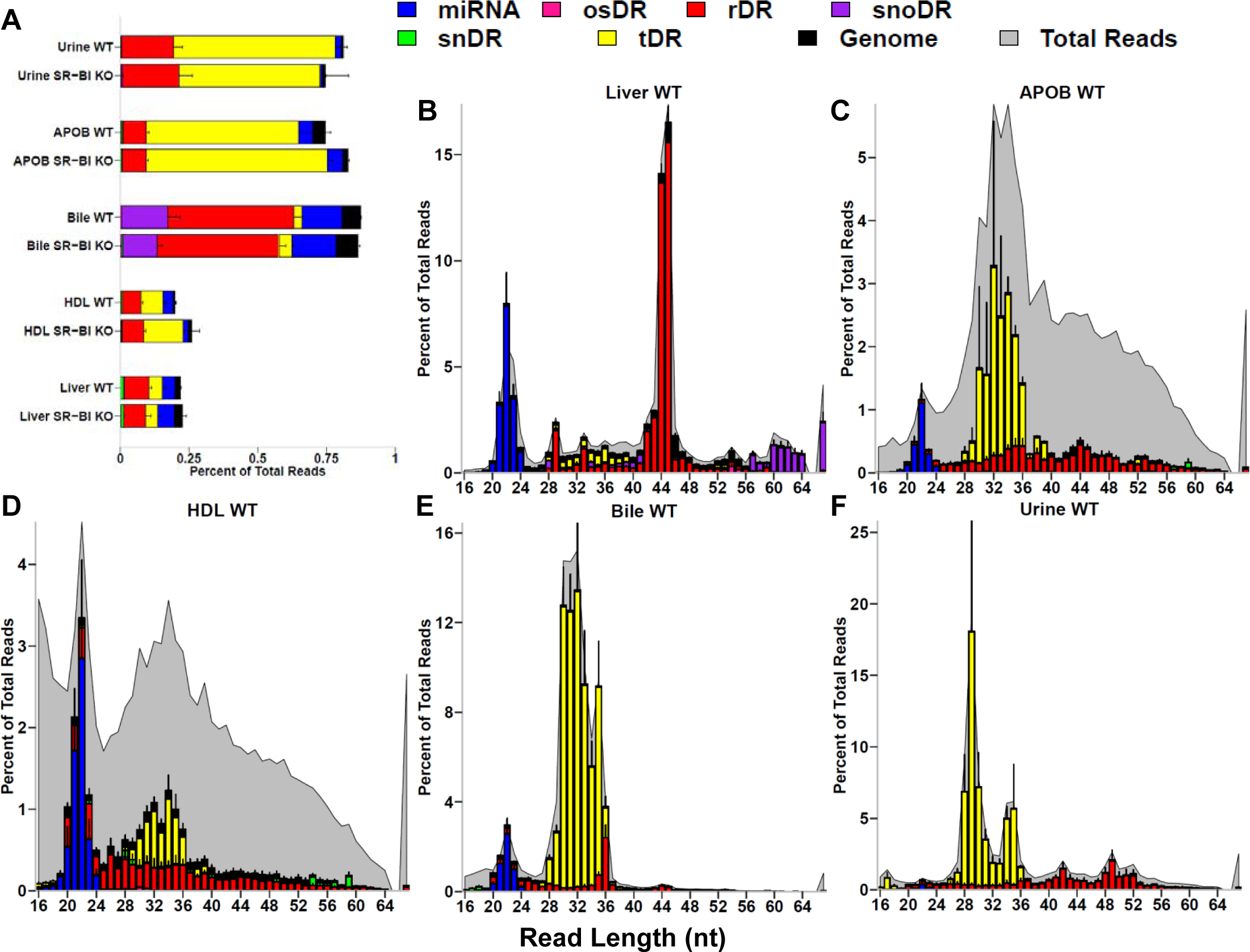
Host sRNAs account for a minor fraction of total reads in lipoprotein sRNA-seq datasets. WT, wild-type mice; SR-BI KO, Scavenger receptor BI Knockout mice (*Scarb1*^*-/-*^). (**A-F**) Results from sRNA-seq analysis. HDL WT, N=7; HDL SR-BI KO N=7; APOB WT, N=7, APOB SR-BI KO N=7; Liver WT, N=7; Liver SR-BI KO, N=7; Bile WT, N=7; Bile SR-BI KO, N=6; Urine WT, N=5; Urine SR-BI KO, N=6. Host tDRs (yellow), rDRs (red), miRNAs (blue), snoDRs (purple), snDRs (green), miscellaneous RNA (pink), and unannotated genome (black). (**A**) Percent of total reads for host sRNA classes. Mean ±S.E.M. (**B-F**) Distribution of read length by host sRNA classes (colors) and total reads (gray), as reported by percent of total reads. Mean ±S.E.M. (**B**) Liver. (**C**) APOB particles. (**D**) HDL. (**E**) Bile. (**F**) Urine.

**Figure 4.**
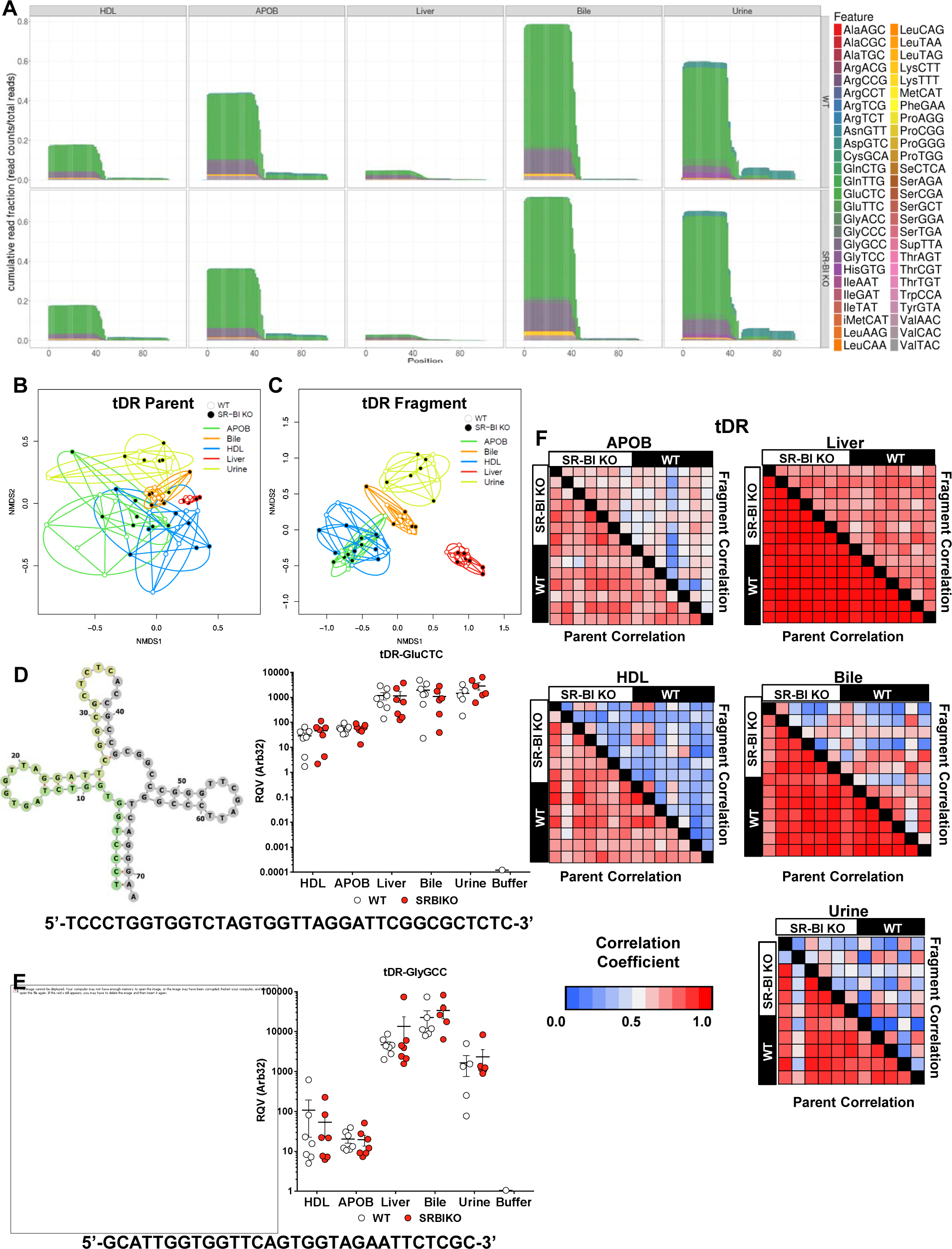
Lipoproteins, bile, and urine contain distinct tDR profiles. WT, wild-type mice; SR-BI KO, Scavenger receptor BI Knockout mice (*Scarb1*^*-/-*^). (**A-C,F**) Results from sRNA-seq analysis. (**A**) Positional coverage maps of tDRs for parent tRNA amino acid anti-codons, as reported as mean cumulative read fractions (read counts / total counts). (**B-C**) Principal Coordinate Analysis (PCoA) of tDR profiles based on (**B**) parent tRNAs and (**C**) individual fragments for samples from WT (empty circles) and SR-BI KO (filled circles) mice. NMDS1, Non-metric multidimensional scaling. (**D-F**) Real-time PCR analysis of candidate tDRs with predicted folding structures and sequences for (**D**) tDR-GluCTC and (**E**) tDR-GlyGCC. WT, white circles; SR-BI KO, red circles. (**F**) Heatmaps of correlation coefficients (Spearman, R) for tRNA parents and individual tDR fragments across samples within each group. HDL WT, N=7; HDL SR-BI KO N=7; APOB WT, N=7, APOB SR-BI KO N=7; Liver WT, N=7; Liver SR-BI KO, N=7; Bile WT, N=7; Bile SR-BI KO, N=6; Urine WT, N=5; Urine SR-BI KO, N=6.

To validate candidate host sRNAs on lipoproteins and in biofluids identified by sRNA-seq, real-time PCR using custom locked-nucleic acid (LNA)-based assays (Exiqon) were completed. For tDRs, both tDR-GluCTC (38 nts in length) and tDR-GlyGCC (32 nts in length) were confirmed to be highly-abundant on HDL and APOB particles, and were not detected in the negative control (buffer) solution used to isolate the lipoproteins (**Figure 4D,E**). Furthermore, two novel snDRs and a candidate sRNA cleaved from a ribozyme (miscRNA) were also detected by PCR on lipoproteins at comparable levels to a previously reported miRNA on lipoproteins (miR-223-3p) (**Figure 4**-**Figure Supplements 5A-D**). Although the general, regional cleavage patterns for specific parent RNAs were consistent for tRNAs (**Figure 4**-**Figure Supplement 1**) and snRNAs (**Figure 4**-**Figure Supplement 6**), and specific candidate sRNAs can be quantified by PCR as single products (based on melting curves), most individual fragments were variable across samples due to slight differences in length or sequences. To more clearly illustrate this point, we performed correlations at both the parent RNA and individual fragment levels within each sRNA class. For tDRs (**Figure 4F**) and other RNA classes (**Figure 4**-**Figure Supplement 7**), we found high correlation between samples at the parent level and poor correlation between samples at the fragment level for HDL, APOB, bile, and urine. For liver samples, high correlation was detected for sRNAs at both the parent and fragment levels (**Figure 4F**, **Figure 4**-**Figure Supplement 7**). These results suggest that, although individual fragments define sRNA classes across groups, further investigation of individual candidate sRNAs (fragments) may be challenging due to variability across samples.

### Lipoproteins are enriched in exogenous sRNAs

Reads aligning to non-human transcripts have previously been detected in human plasma samples^1^; however, it is unknown which carriers transport non-host sRNAs in host circulation. To determine if lipoproteins carry exogenous bacterial and fungal sRNAs, reads >20 nts in length that failed to map to the host (mouse) genome were aligned in parallel to A.) Annotated non-host transcripts curated in GtRNAdb (tRNA), SILVA (rRNA), and miRBase (miRNA) databases, and B.) Genomes of bacteria and fungi of the microbiome (human microbiome, HMB) or environment (ENV) (**Figure 1**). To identify exogenous miRNAs (xenomiRs), reads were aligned (perfect match only) to non-host mature miRNA sequences (miRBase.org); however, only a few xenomiRs were detected within our datasets and overall contributions to each profile were minimal (**Figure 5**-**Source Data 1**). To determine the levels of exogenous tDRs on lipoproteins, non-host reads were aligned to parent tRNAs curated in the GtRNAdb library. Both HDL and APOB particles were found to transport a diverse set of exogenous tDRs across multiple kingdoms, which accounted for approximately 2.5% of sRNAs (total reads) circulating on each lipoprotein class (**Figure 5A**, **Figure 5**-**Source Data 2**). Bacterial tDRs were the most represented taxon, and the most abundant bacterial species were Pseudomonas fluorescens, Pseudomonas aeruginosa, and Acinetobacter baumanni (**Figure 5**-**Figure Supplements 1A,B**). The parent tRNAs (based on amino acid anticodons) with the highest normalized read counts were fMetCAT, GluTTC, AspGTG, and AsnGTT (**Figure 5**-**Figure Supplement 1C**). In contrast to host tDRs that predominantly aligned to the 5’ halves of parent tRNAs (**Figure 4A**, **Figure 4**-**Figure Supplement 1**), positional coverage analysis demonstrated that bacterial tDRs aligned across the entirety of tRNA transcripts (**Figure 5B**, **Figure 5**-**Figure Supplements 2 and 3**). To identify exogenous rDRs on lipoproteins, non-host reads were separately aligned to known rRNA transcripts curated in the SILVA database, and remarkably, non-host rDRs accounted for approximately 25% of total reads in each of the HDL and APOB datasets (**Figure 5C**, **Figure 5**-**Source Data 3**). Although rDRs from every taxonomical kingdom were present on lipoproteins, bacterial rDRs were the most abundant (**Figure 5C**, **Figure 5**-**Figure Supplement 4, Source Data 3**). The overall content of non-host sRNAs on HDL and APOB particles were similar; however, HDL were found to be enriched for shorter length non-host tDRs and rDRs compared to APOB particles (**Figures 5D,E**). Collectively, these results suggest that lipoproteins transport exogenous tDR and rDRs, most of which are likely bacterial in origin.

**Figure 5.**
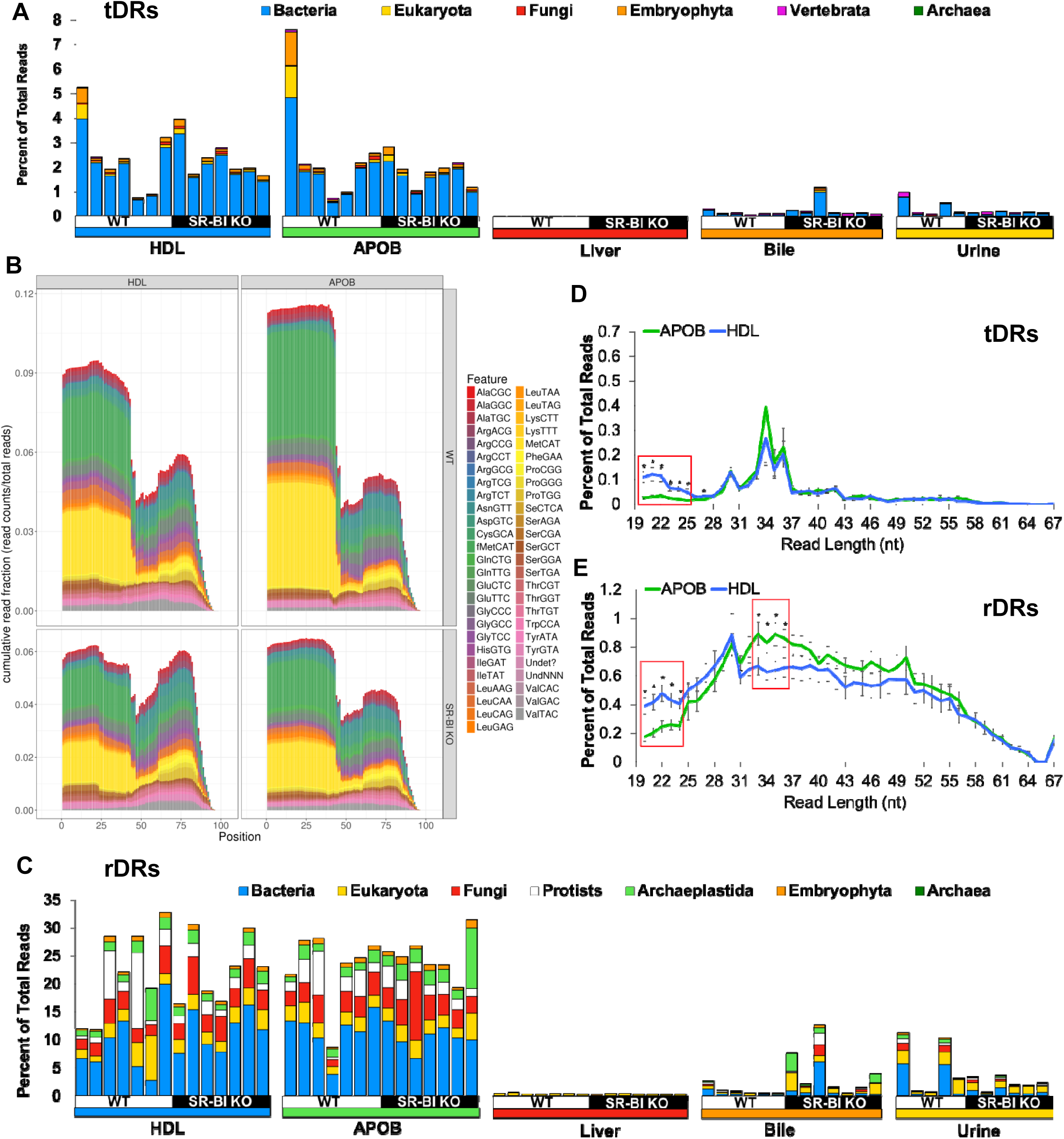
Lipoproteins transport exogenous non-host tDRs and rDRs. WT, wild-type mice; SR-BI KO, Scavenger receptor BI Knockout mice (*Scarb1*^*-/-*^). (**A**) Stacked bar plots of non-host tDRs aligned to parent tRNAs across kingdoms and higher organizations – bacteria, blue; eukaryota, yellow; fungi, red; embryophyta, orange; vertebrata, purple; archaea, green – as reported as percent of total reads. (**B**) Positional coverage maps of non-host tDRs for parent tRNA amino acid anti-codons, as reported as mean cumulative read fractions (read counts / total counts) for HDL and APOB particles. (**C**) Stacked bar plots of non-host rDRs aligned to parent rRNAs across kingdoms and higher organizations – bacteria, yellow; eukaryota, red; fungi, white; protists, purple; archaeplastida, dark blue; embryophyta, light blue; archaea, green – as reported as percent of total reads. (**D-F**) Distribution of read lengths, as reported as percent of total reads, for all non-host (**D**) tDRs and (**F**) rDRs. Two-tailed Student’s t-tests. *p<0.05. HDL WT, N=7; HDL SR-BI KO N=7; APOB WT, N=7, APOB SR-BI KO N=7; Liver WT, N=7; Liver SR-BI KO, N=7; Bile WT, N=7; Bile SR-BI KO, N=6; Urine WT, N=5; Urine SR-BI KO, N=6.

Aligning reads to transcripts in databases is biased in that only known (annotated) RNAs are queried, and thus, limits the power of discovery in sRNA-seq datasets. To comprehensively analyze exogenous sRNAs, non-host reads were also aligned to bacterial genomes within the NIH HMB Project (hmpdacc.org). The HMB database currently holds 3,055 genomes, many of which are closely related; therefore, to address potential multi-mapping issues, we collapsed these species into 206 representative genomes that spanned 11 phyla and accounted for every genera within the HMB. Alignment of non-host reads to HMB genomes identified many bacterial sRNAs on lipoproteins and in biofluids, reported as summarized genome read counts per million total reads (RPM) (**Figure 6-Figure Supplement 1A-B, Source Data 1**). To perform taxonomical analyses of lipoprotein-associated bacterial sRNAs, circular tree maps were generated. As shown by concentric rings in the tree maps, the vast majority of both HDL and APOB bacterial reads mapped to the Proteobacteria phylum (green), followed by the Actinobacteria (blue) and Firmicutes (yellow) phylums (**Figure 6A**, **Figure 6**-**Figure Supplement 2A**). Within the Proteobacteria phylum, the majority of reads aligned to the Gammaproteobacteria class, particularly the orders of Pseudomonadales and Enterobacteriales. Among individual genera (inner-most circles), counts for the genus Pseudomonas (Proteobacteria phylum) were consistently the high, as were Micrococcus (Actinobacteria phylum) (**Figure 6A**, **Figure 6**-**Figure Supplement 2A**).

**Figure 6.**
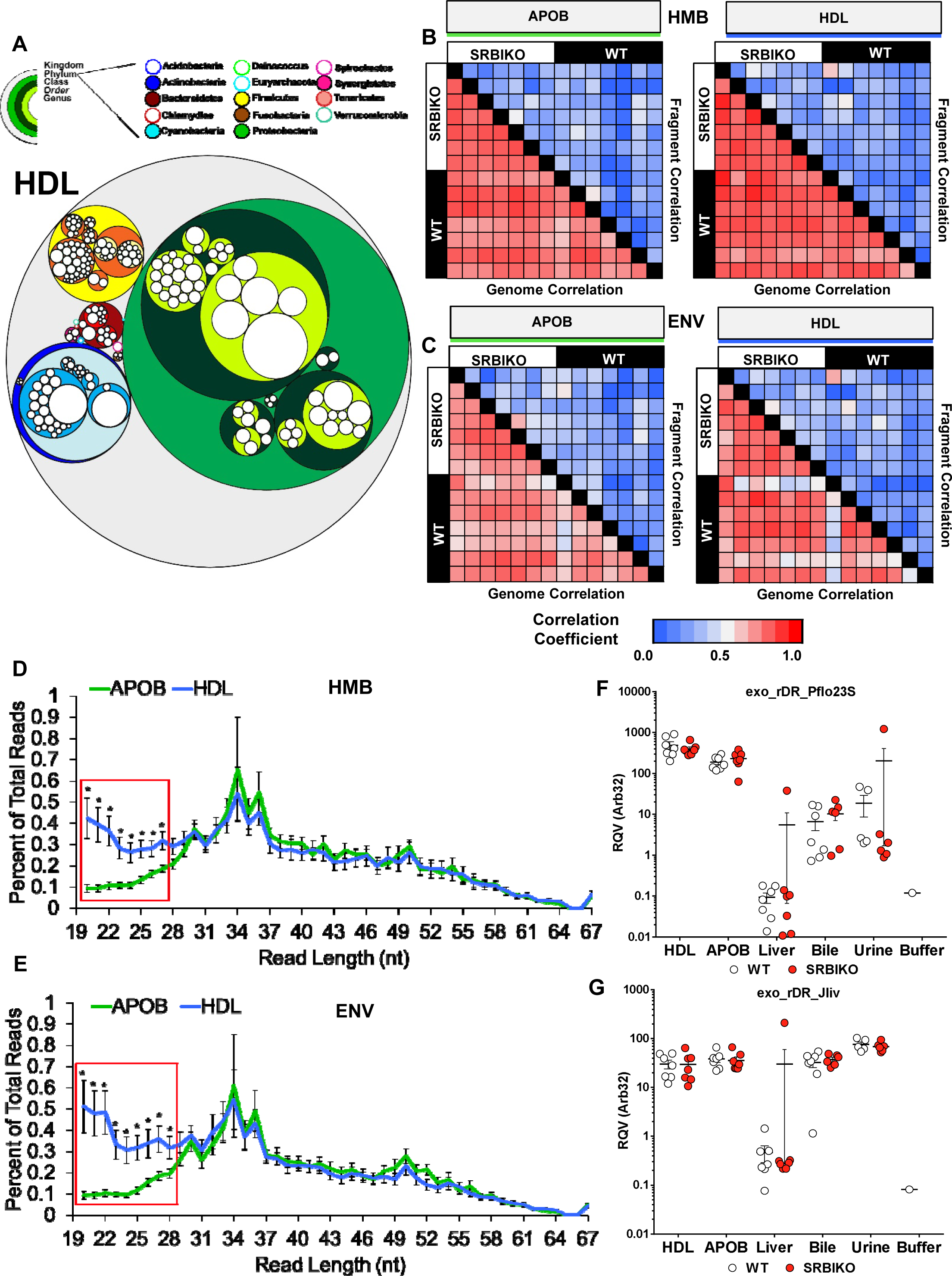
Lipoproteins are enriched for sRNAs derived from proteobacteria in the microbiome and environment. WT, wild-type mice; SR-BI KO, Scavenger receptor BI Knockout mice (*Scarb1*^*-/-*^). (**A**) Circular tree maps for non-host bacterial sRNAs on HDL from WT mice, as organized by taxonomy – proteobacteria, green; actinobacteria, blue; firmicutes, yellow; bacteroidetes, red. Diameter is proportional to the mean number of reads at the genome level (counts). (**B**-**C**) Heatmaps of correlation coefficients (Spearman, R) for non-host sRNAs (on HDL and APOB particles) for bacterial genomes and individual bacterial fragments across samples grouped by (**B**) human microbiome (HMB) and (**C**) environment (ENV) species. (**D-E**) Distribution of read lengths, as reported as percent of total reads, for non-host bacterial sRNAs grouped by (**D**) HMB and (**E**) ENV species. Two-tailed Student’s t-tests. *p<0.05. (**F**-**G**) Real-time PCR analysis of candidate non-host bacterial sRNAs for (**F**) exogenous rDR Pseudomonas fluorescens 23S (exo_rDR_Pflo23S) and (**G**) exogenous rDR Janthinobacterium lividum 23S (exo_rDR_Jliv). WT, white circles; SR-BI KO, red circles. HDL WT, N=7; HDL SR-BI KO N=7; APOB WT, N=7, APOB SR-BI KO N=7; Liver WT, N=7; Liver SR-BI KO, N=7; Bile WT, N=7; Bile SR-BI KO, N=6; Urine WT, N=5; Urine SR-BI KO, N=6.

Most interestingly, many reads that aligned to bacterial rRNA transcripts failed to align to the HMB genomes, thus suggesting that some sRNAs may originate from bacteria not presently curated in the HMB database. Using BLASTn (NCBI), many highly abundant reads were perfect matches to bacterial genomes of environmental bacterial species of soil and water, but could be associated with opportunistic infections. Therefore, to increase our non-host coverage, 167 additional bacterial genomes representing non-redundant genera of 8 taxonomical phyla were added, termed here as environmental bacteria (ENV). The ENV species with the highest normalized genome counts for WT lipoproteins were Pseudomonas fluorescens, Pseudomonas putida, Propionibacterium acnes, and Stenotrophomonas maltophilia (**Figure 6**-**Figure Supplements 1C,D, and 2B,C, Source Data 2**). Although many non-host reads aligned to both HMB and ENV genomes, a majority of all non-host bacterial reads could be assigned exclusively to only one database, suggesting a complex origin for bacterial sRNAs on circulating lipoproteins (**Figure 6**-**Figure Supplements 3**). In addition to bacterial sRNAs, we also identified fungal sRNAs on lipoproteins, and the highest normalized genome counts for fungal species on WT HDL were Fusarium oxysporum, Histoplasma capsulatum, Cryptococcus neoformans (**Figure 6**-**Figure Supplements 4A,B, Source Data 3**).

To assess bacterial sRNA profiles across samples, non-host sRNAs (HMB and ENV) on lipoproteins were correlated between samples. For both databases, we identified high correlations between samples at the genome level and low correlations at the fragment level (**Figures 6B,C**). These data suggest that similar bacteria are contributing sRNAs to circulating lipoproteins across all mice. Nevertheless, these bacteria may contribute different sRNAs (sequences) to HDL and APOB particles in different mice or the processing of bacterial sRNAs before and/or during HDL and APOB trafficking is differentially regulated. A key difference between HDL and APOB bacterial sRNAs was length, as HDL were enriched for shorter sRNAs than APOB particles; this pattern was evident for both HMB and ENV sRNAs (**Figures 6D,E**). A similar trend was observed for reads mapping to fungal genomes (**Figure 6**-**Figure Supplement 5**). To determine if HDL and APOB particles transport different exogenous (non-host) sRNA signatures, PCoA and PERMANOVA analyses were completed. At the genome level, the HDL and APOB particles were indistinguishable for HMB and ENV bacteria (**Figure 6**-**Figure Supplement 6A,B**); however, HDL and APOB profiles clustered separately at the fragment level and HDL and APOB profiles were distinct (F=1.7, p=0.048) for ENV bacterial sRNAs by PERMANOVA (**Figure 6**-**Figure Supplement 6C,D, Source Data 4**).

The lack of strong correlation at the fragment level for non-host sRNAs is likely due to differences in read lengths and sequences (e.g. terminal nts) for similar reads due to imprecise processing of parent RNAs, and thus variable read counts across samples. These observations present unique challenges to study individual sRNAs for biological function; however, many candidate sRNAs do exist within the very large pool of non-host reads. Using real-time PCR, we quantify candidate bacterial sRNAs on lipoproteins, and confirmed that HDL and APOB particles transport a 22 nt rDR (5’-AGAGAACUCGGGUGAAGGAACU-3’) likely from bacteria of the Proteobacteria phylum (**Figure 6F**, **Figure 6**-**Figure Supplement 7**). Likewise, HDL and APOB were also found to transport another rDR of the Proteobacteria phylum, likely from the order of Burkholderiales (33 nts, 5’-GACCAGGACGUUGAUAGGCUGGGUGUGGAAGUG-3’) (**Figure 6G**, **Figure 6**-**Figure Supplement 8**). In addition to bacterial sRNAs, real-time PCR was also used to confirm that lipoproteins transport a fungal rDR from the Verticillium genus (21 nts 5’-UGGGUGUGACGGGGAAGCAGG-3’) (**Figure 6**-**Figure Supplement 9**). These results suggest that HDL and APOB transport non-host sRNAs derived from bacterial and fungal sources in the microbiome and environment. Bile and urine samples also contained non-host sRNAs, albeit a lesser fraction of total reads. Collectively, these observations support the need to include non-host sRNAs in the analysis of sRNA-seq data.

### Lipoproteins are defined by their most abundant sRNAs

To determine which RNA class and species contribute to the most abundant sRNAs in each sample type, the top 100 ranked reads for each sample were filtered and redundant reads were removed for each group. For liver and bile samples, the top ranked reads were predominantly host sRNAs (**Figure 7A,B**). On the contrary, the top most abundant reads on lipoproteins were comprised of both host and non-host sRNAs (**Figures 7C,D**). The top ranked reads in urine samples were found to be largely host sRNAs (e.g. tDRs); however, many links to exogenous bacterial sRNAs were identified (**Figure 7E**). Although our host and non-host analyses were thorough, many of the top ranked sequences remained unidentified. Therefore, we sought to further analyze sRNA profiles using a class-independent strategy, in which we focused on only the most abundant reads for each group. To assess the similarity of profiles between groups for the top ranked sRNAs, hierarchical clustering and correlations were performed, and lipoproteins displayed modest correlations with other groups and clustered separately from livers, bile, and urine (**Figure 7**-**Figure Supplement 1**). These observations were confirmed by PCoA, as lipoprotein samples overlapped and clustered together, separately from bile, urine and liver samples (**Figure 7F**). PERMANOVA analysis found that every group was significantly distinct from each other based on the most abundant sRNAs (**Figure 7**-**Source Data 1**). These results suggest that each sample type can be defined by their most abundant sRNAs independent of parent RNA class or contributing host or non-host species which is highly appropriate for the study of heterogeneous pools of exRNAs.

**Figure 7.**
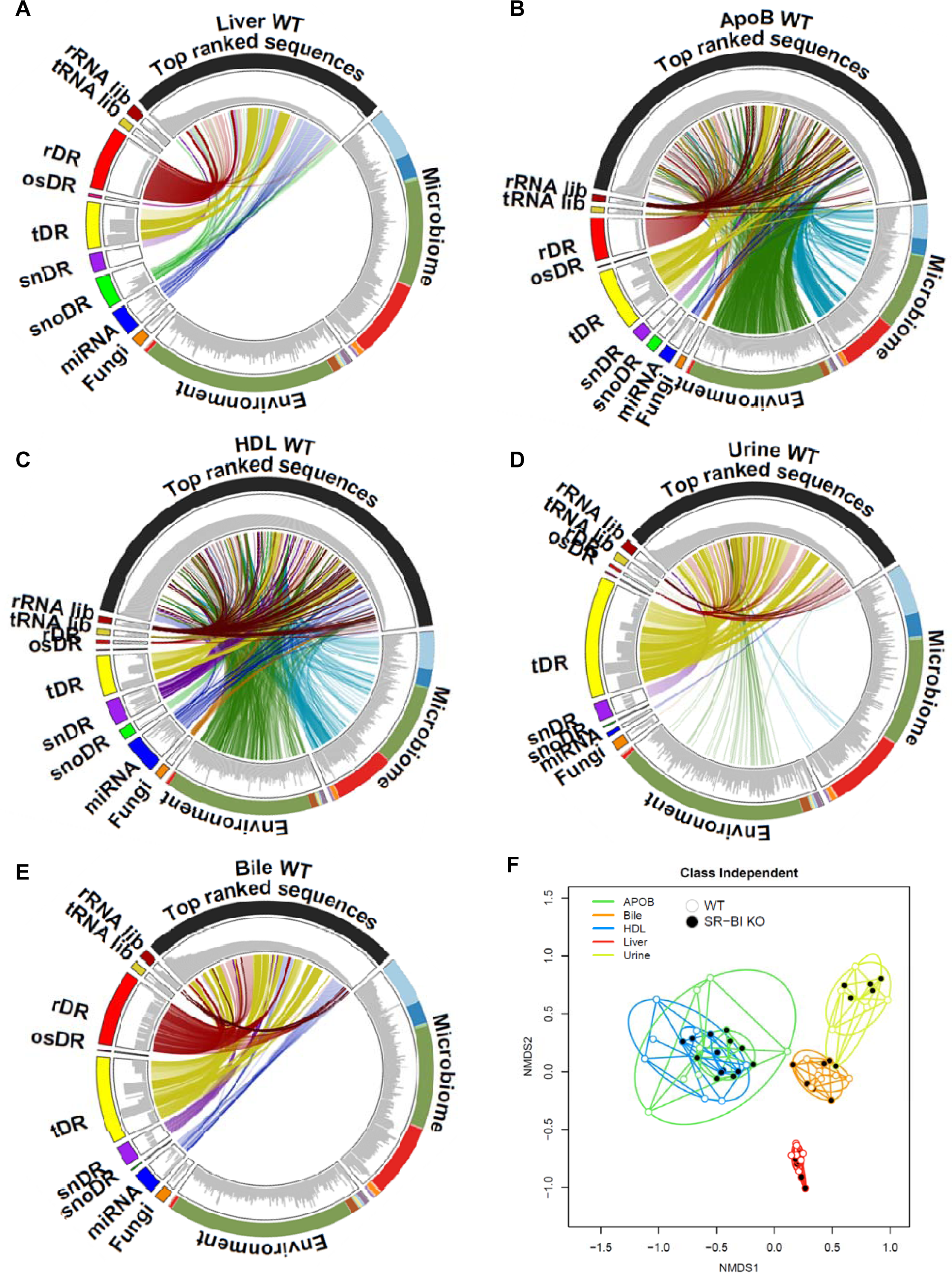
The most abundant sRNAs on lipoproteins are bacterial rDRs. (**A-E**) Circos plots linking the most abundant (top 100) sequences to assigned groups for non-host libraries (rRNA lib, tRNA lib), host sRNAs (rDR, osRNA, tDRs, snDRs, snoDRs, miRNAs) and non-host genomes (fungi, environment, and microbiome) for (**A**) liver, (**B**) bile, (**C**) APOB, (**D**) HDL, and (**E**) urine. (**F**) Principal Coordinate Analysis (PCoA) of sRNA profiles based on class-independent analyses. Wild-type mice, WT (open circles); Scavenger receptor BI Knockout mice (*Scarb1*^*-/-*^), SR-BI KO (filled circles). HDL WT, N=7; HDL SR-BI KO N=7; APOB WT, N=7, APOB SR-BI KO N=7; Liver WT, N=7; Liver SR-BI KO, N=7; Bile WT, N=7; Bile SR-BI KO, N=6; Urine WT, N=5; Urine SR-BI KO, N=6.

### Advances in sRNA-seq analysis

To compare the TIGER pipeline to other sRNA-seq analysis software, APOB, HDL, and liver samples from WT mice were analyzed by Chimira^8^, Oasis^14^, ExceRpt^25^, and miRge^13^ software (**Figure 8**-**Source Data 1**). Although the pipelines are designed for different outputs, each can quantify host miRNAs for which we used to compare analyses, and we found that all the pipelines were comparable in their ability to quantify host canonical miRNAs for different sample types and the pipelines were highly correlated for miRNAs (**Figure 8**-**Figure Supplement 1A,B**). Most available software for sRNA-seq data analysis are restricted to miRNAs or endogenous (host) sRNAs, including Chimira, Oasis, and miRge (**Figure 8**-**Source Data 1**). This approach may be suitable for liver samples (red circles), as demonstrated by ternary plots, but HDL (blue circles) and APOB (green circles) samples remain largely unexplained (**Figure 8A**). Incorporation of both endogenous and exogenous sRNAs, a key feature of the TIGER pipeline, is essential to studying lipoprotein sRNAs as this strategy accounts for substantially more reads in the datasets, as depicted by the left shifts of blue and green circles in the ternary plots (**Figure 8B**, **Figure 8**-**Source Data 2**). A key metric for comparing pipelines is the amount of (useable) information extracted from the data by the software, i.e. the percent of assigned quality reads. Remarkably, the TIGER pipeline accounted for 87.95% bile, 87.9% of liver, 85.3% urine, 71.5% HDL, and 62.2% APOB reads in WT mice (**Figure 8C**, **Figure 8**-**Source Data 3**). In comparison to other pipelines, the TIGER pipeline accounted for significantly more reads in lipoprotein datasets, and significantly more reads than Chimira, Oasis, and ExceRpt for liver datasets which are largely host sRNAs (**Figures 8C,D**). After the TIGER pipeline performs the non-host read analyses, the top ranked most abundant sequences of the unexplained reads that remain were searched using BLASTn (**Figure 1**). Collectively, the TIGER pipeline provides an opportunity to analyze sRNA-seq with increased depth and detail which is particularly suited for analysis of exRNA and sRNAs on lipoproteins.

**Figure 8.**
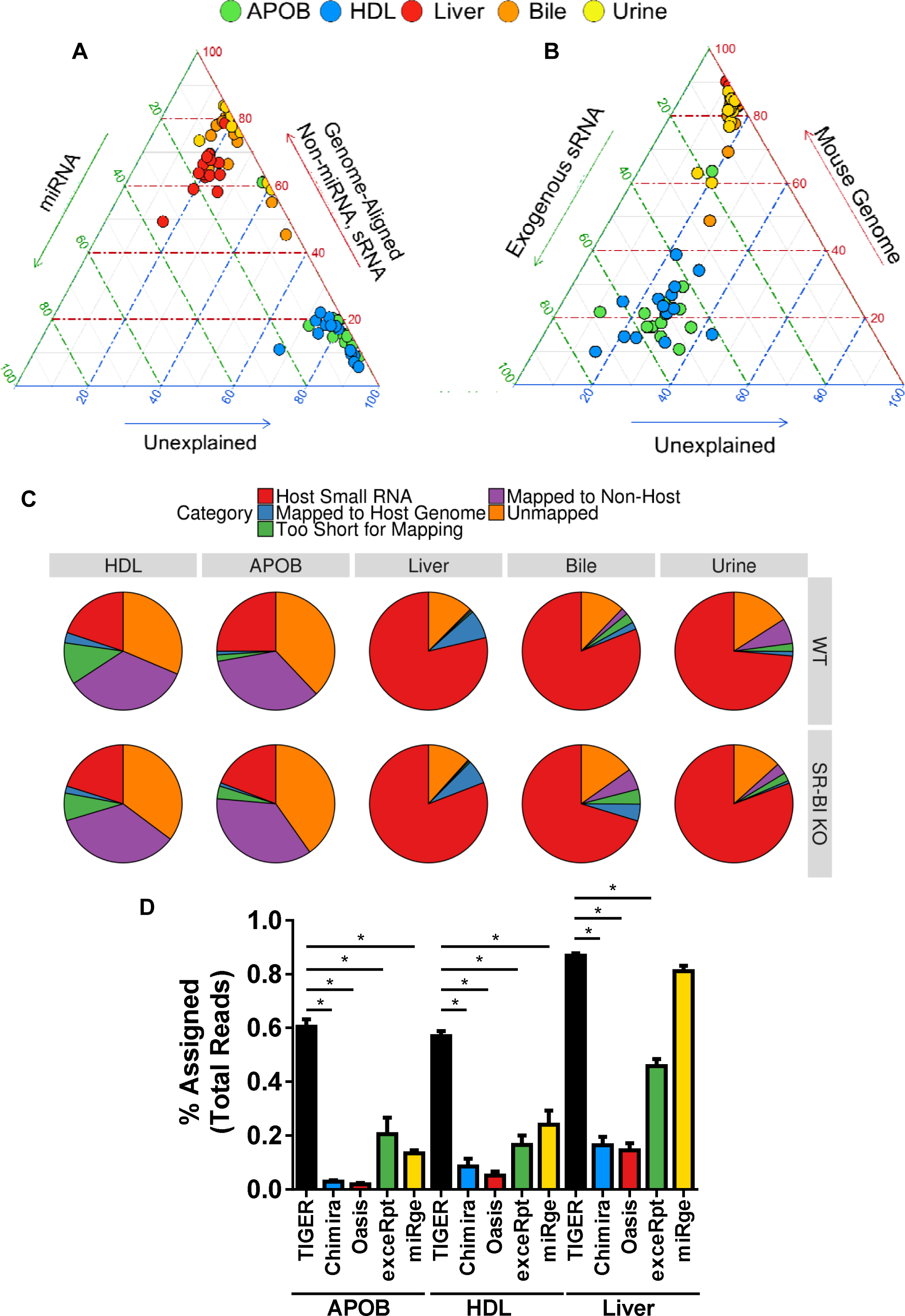
TIGER analysis pipeline accounts for significantly more reads than other software for lipoprotein sRNA-seq data. (**A-B**) Ternary plots of sRNA profiles for all samples displayed as (**A**) percent unexplained (blue), miRNAs (green), and non-miRNA host sRNAs (red); (**B**) percent unexplained (blue), exogenous sRNAs (green), and host genome (red). WT, wild-type mice; SR-BI KO, Scavenger receptor BI Knockout mice (*Scarb1*^*-/-*^). (**C**) Pie charts illustrating the mean fraction of reads assigned to host sRNA (red), host genome (blue), non-host (purple), too short for mapping (green), and unmapped (orange). HDL WT, N=7; HDL SR-BI KO N=7; APOB WT, N=7, APOB SR-BI KO N=7; Liver WT, N=7; Liver SR-BI KO, N=7; Bile WT, N=7; Bile SR-BI KO, N=6; Urine WT, N=5; Urine SR-BI KO, N=6. (**D**) Comparisons of sRNA-seq data analysis pipelines, as reported as percent assigned per total reads for TIGER (black), Chimira (blue), Oasis (red), ExceRpt (green), and miRge (yellow) for HDL, APOB, and liver samples from WT mice. HDL WT, N=7; APOB WT, N=7, Liver WT, N=7. Mann-Whitney non-parametric tests. *p<0.05.

### SR-BI Regulation of Lipoprotein sRNAs

SR-BI is highly-expressed in the liver and plays a fundamental role in reverse cholesterol transport mediating hepatic uptake of HDL-cholesteryl esters and biliary cholesterol secretion^26–28^. Loss-of-function variants in human *SCARB1* (SR-BI) were associated with increased in circulating HDL-C levels^29^. Likewise, *Scarb1* mutations in mice also resulted in increased HDL-C levels^30^. We have previously reported that HDL-delivery of miRNAs to hepatocytes *in vitro* requires SR-BI^17^. Based on these observations, we hypothesized that SR-BI may regulate sRNA levels on lipoproteins as well as miRNAs in liver and bile. To quantify the impact of SRBI-deficiency on exRNAs *in vivo*, host sRNAs were compared at both the parent and fragment levels. For miRNAs, loss of SR-BI in mice did not alter miRNA content in liver, urine, bile, or APOB particles at the parent level, and only one miRNA (mmu-miR-143-3p, 0.199-fold, adjp= 0.00042) was significantly altered in SR-BI KO mice compared to WT mice (**Figure 9A**, **Figure 9**-**Source Data 1**). We also identified a limited number of significantly altered non-miRNA host sRNAs at the parent level in SR-BI KO mice compared to WT mice (**Figure 9A**, **Figure 9**-**Figure Supplements 1**, **Figure 9**-**Source Data 1**). SR-BI may regulate urinary miRNA NTAs as SR-BI KO mice were found to have a significant increase in urinary miRNA NTAs (p<0.001) compared to WT mice (**Figure 2**-**Figure Supplement 1C**). Moreover, we found a significant (p=0.0021) change in NTA-A/U ratios in urine from SR-BI KO mice compared to WT mice, as urine samples from WT mice were enriched for poly-adenylated miRNAs (NTA-A) and samples from SRBI KO mice were enriched for poly-uridylated miRNAs (NTA-U) (**Figure 2F**).

**Figure 9.**
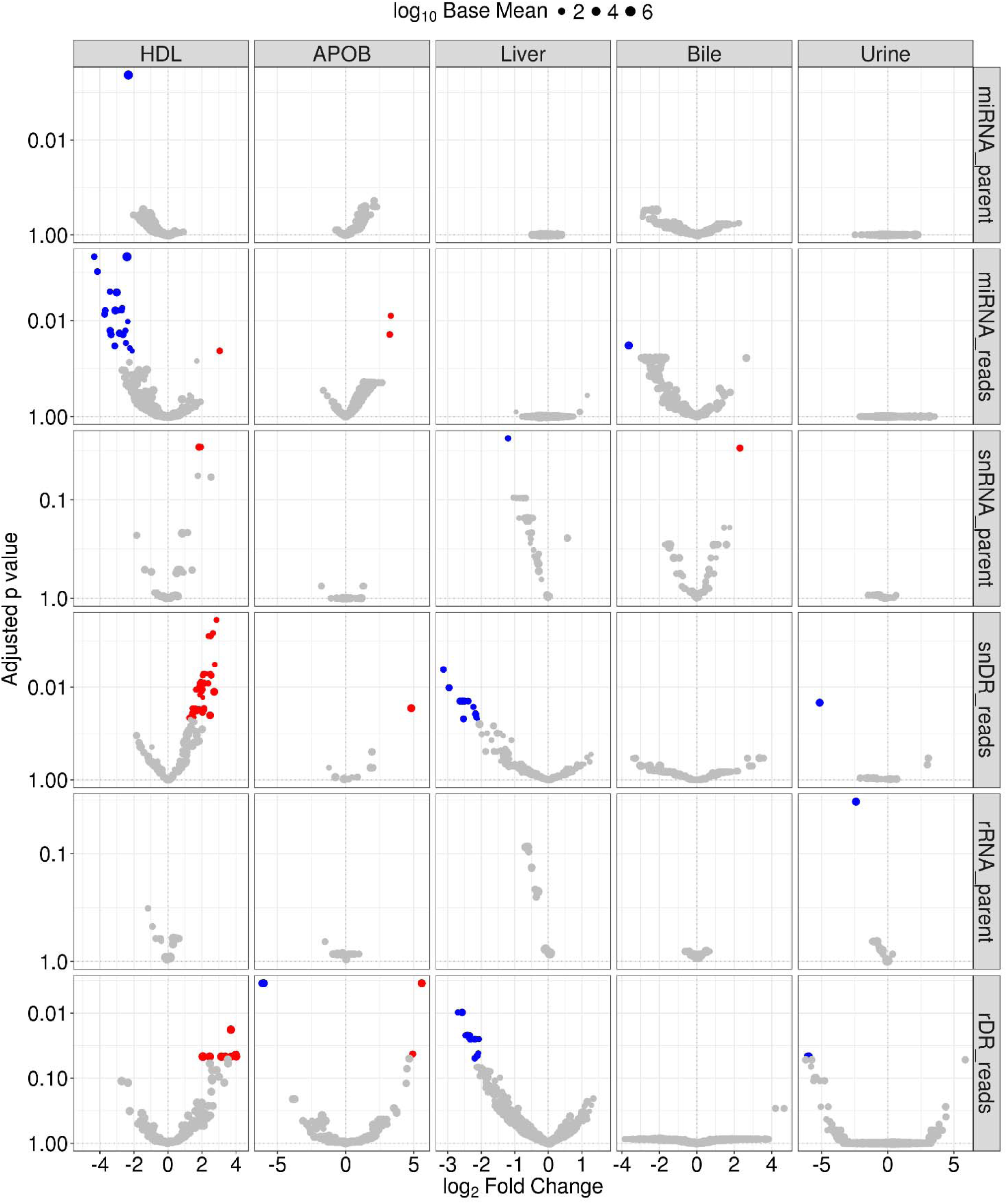
SR-BI regulates HDL-sRNAs at the individual fragment level, not parent level. Differential expression analysis by DEseq2. Volcano plots of demonstrating significant (adjusted p>0.05) differential (>1.5-absolute fold change) abundances for miRNAs, snDRs, and rDRs at the parent and individual fragment levels - red, increased; blue, decreased. HDL WT, N=7; HDL SR-BI KO N=7; APOB WT, N=7, APOB SR-BI KO N=7; Liver WT, N=7; Liver SR-BI KO, N=7; Bile WT, N=7; Bile SR-BI KO, N=6; Urine WT, N=5; Urine SR-BI KO, N=6.

The impact of SR-BI-deficiency on lipoprotein sRNAs may not be evident by grouping individual sRNAs by their likely parent RNAs and differential expression analyses of individual fragments for each RNA class may be necessary, as the expression of many individual fragments were found to be significantly altered in SR-BI KO mice compared to WT mice and distinct patterns were detected (**Figure 9**). For example, SR-BI-deficiency resulted in a significant decrease to 21 individual miRNA sequences (**Figure 9**, **Figure 9**-**Souce Data 2**). Conversely, we found 57 snDR and 8 rDR fragments that were significantly increased on HDL (**Figure 9**-**Souce Data 2**). In livers from SR-BI KO mice, we found 14 snDRs and 16 rDRs that were significantly increased at the fragment level, although these were not identical sequences to fragments found to be decreased on HDL for these classes (**Figure 9**, **Figure 9**-**Souce Data 2**). These results suggest that SR-BI may play a limited role in regulating sRNAs circulating on HDL and in livers. Nevertheless, these results strongly support the need to analyze host sRNAs not just at the parent level, but also the fragment level, as potentially critical observations may be lost in the grouping of similar sequences for parent analysis.

Although bacteria may regulate SR-BI expression^31^, SR-BI regulation of the gut microbiome is unclear, and the role of SR-BI in regulating circulating non-host bacterial sRNAs on lipoproteins is completely unknown. To determine if SR-BI contributes to exogenous sRNAs on lipoproteins and in biofluids, differential expression analysis was performed at both the genome and fragment levels. Only one bacterial species was found to be significantly altered between SR-BI KO and WT mice, decreased Streptomyces in urine, as determined by genome counts (**Figure 9**-**Figure Supplement 2A**, **Figure 9**-**Source Data 3**). At the individual fragment level, only 3 bacterial sRNAs were significantly altered by SR-BI-deficiency; one each in APOB, bile, and urine samples (**Figure 9**-**Figure Supplement 2B**, **Figure 9**-**Source Data 4**). These results suggest that SR-BI does not likely regulate non-host bacterial sRNAs on lipoproteins or in biofluids. Conversely, we found a significant increase in all fungal genome counts in SR-BI KO mice compared to WT mice (**Figure 9**-**Figure Supplement 2A**, **Figure 9**-**Source Data 3**). These observations were not likely the result of a few dominant reads shared across all fungal genomes as we failed to find any individual fungal sRNAs that were significantly affected by the loss of SR-BI (**Figure 9**-**Figure Supplement 2B**). To determine if SR-BI-deficiency in mice results in changes to the most abundant sequences in each group, independent of RNA class or genotype, differential expression analysis was performed for the top 100 reads filtered in the class-independent analysis. Nonetheless, we only found changes to the most abundant reads on lipoproteins in SR-BI KO mice compared to WT mice (**Figure 9**-**Figure Supplement 3**, **Figure 9**-**Source Data 5**).

## Discussion

High-throughput sequencing of sRNAs has revealed a complex landscape of various types of sRNAs in cells and extracellular fluids, many of which have not been studied. Currently, there is a great need for tools that can extract many types of sRNAs and their distinct features from sequencing datasets. Here, we used sRNA-seq and TIGER to profile most sRNA classes on HDL and APOB particles and compared these profiles to liver, bile, and urine. Using this approach, we found that HDL and APOB particles transport a wide-variety of host sRNAs, including tDRs, rDRs, snDRs, and many other miscRNAs. Moreover, we found that exRNAs on lipoproteins harbored unique features, such as, enrichment of poly-uridylation NTA events on miRNAs and discrete length distributions for HDL and APOB particles. Furthermore, lipoproteins were found to transport a multitude of non-host sRNAs likely derived from bacterial and fungal species of the microbiome and environment. Many of these non-host sRNAs were found to be likely processed from parent tRNAs and rRNAs. Using TIGER, we were also able to define each sample type by their most abundant sRNAs independent of class or species, which is particularly suited for the study of exRNA. Furthermore, the TIGER pipeline allows for the quantification and differential expression analysis of sRNAs at both the parent and fragment levels. This strategy allowed our determination that SR-BI has a limited role in regulating cellular and extracellular sRNAs, which would not have been feasible with other analysis strategies that focus solely on the parent RNA organization. Overall, this study demonstrates the power of expanding sRNA-seq analysis beyond canonical miRNAs and exploring the full breadth of host and non-host sRNAs in every dataset.

Although many researchers are using high-throughput sequencing to quantify sRNAs, many investigators do not take advantage of the enormous amount of information contained within sRNA-seq datasets. The mammalian transcriptome is immensely diverse and complex, and thus, requires new analytical tools and novel strategies to address the many distinct features of different sRNA classes and contributing species^10, 12, 32^. TIGER is designed to incorporate both host and non-host sRNA analysis into a modular design that allows for custom prioritization and parallel alignments to both genomes and transcripts (libraries), and organizes data at the parent RNA, fragment, and class-independent levels. The seven modules include preprocessing, host genome and database, non-host library, non-host genome, class-independent, summary, and unmapped. For host miRNAs, we expanded miRNA analysis to include 5’ and 3’ terminal isomiRs and 3’ NTAs. Furthermore, we extended our analysis of annotated host sRNAs to include tDRs, rDRs, snDRs, snoDRs, lncDRs, and many other less studied classes, e.g. yDRs. A key feature of TIGER is the alignment strategy for host tDRs and rDRs which includes mapping to the host genome and mature transcripts in corresponding databases, which overcomes challenges posed by introns^24, 33^. Another key advance in our pipeline is the parallel analysis of host sRNAs at the parent and individual fragment levels. Organization of sRNAs at the parent level allows for categorical analysis and positional coverage alignments which provides information on parent RNA processing (cleavage). Conversely, analysis of sRNAs at the individual sequence (fragment) level aids biomarker discovery and is critical to determining biological functions. Collectively, these features represent a substantial advance for the analysis of endogenous host sRNAs across all types of samples.

A critical difference between cellular RNA and exRNA profiles is the presence of non-host sRNAs present in exRNA samples^1, 34, 35^. ExRNAs hold great potential as disease biomarkers, indicators of specific cell phenotypes and damage, intercellular communication signals, and drug targets for future therapies^36–38^. Current sRNA-seq analysis pipelines are not particularly suitable for the study of exRNAs as many are restricted to only canonical miRNAs, or a limited number of host sRNAs, and lack analysis of non-host sRNAs, which will likely be a major focus of future investigations. Based on a previous study reporting that bacterial sRNAs are present in human plasma, TIGER was designed to identify exogenous bacterial and fungal sRNAs. Strikingly, we found that the majority of sRNAs on HDL and APOB particles are likely from bacteria present in the microbiome and environment. These non-host sRNAs are not likely contamination products due to several observations. First, we were not able to detect candidate bacterial sRNAs in control buffer used to isolate the lipoproteins by real-time PCR. Moreover, reads aligning to bacterial and fungal genomes were not likely contamination of reagents used for sequencing preparation as most of these reads were not present in liver datasets. Next, we found very low correlation between lipoprotein samples for non-host bacterial and fungal sRNAs suggesting that there was not a common source of bacterial or fungal RNA in the preparation reagents. In addition, we found that bacterial and fungal sRNAs on HDL were enriched for short length sRNAs as compared to APOB particles, a pattern that was also observed for host sRNAs, thus supporting a common mechanism of loading or association for sRNAs that is different for HDL and APOB particles. Moreover, we found that non-host bacterial sRNA profiles were distinct for HDL and APOB at the fragment level, as demonstrated by PCoA and PERMANOVA. Collectively, these results strongly support that HDL and APOB particles transport distinct sets of non-host sRNAs that are not likely due to bacterial and fungal contamination or foreign RNA in reagents or the research environment.

The inclusion of non-host reads in our analysis greatly increased our ability to account for reads in lipoprotein datasets. Nevertheless, there are many exogenous sRNAs that could be neither processed from annotated transcripts in databases nor originate from species currently represented in the HMB project. Therefore, another key feature of the TIGER pipeline is the ability to analyze data independent of species identification or library annotation. As such, class-independent analysis extracts more data and eliminates a potential barrier to the discovery of biomarkers and intercellular communication signals. Notably, class-independent analysis of exRNAs captures sRNA sequence, length, and abundance which are the important defining characteristics of biomarkers in extracellular fluids and bioactivity in recipient cells. The TIGER pipeline also advances sRNA-seq analysis through the incorporation of high-end comparative analyses and data visualizations, including PCoA, PERMANOVA, hierarchical clustering and correlations, positional coverage maps, circular tree maps, circos linkage maps, and ternary plots. The TIGER pipeline addresses many issues in sRNA-seq analysis; however, we have identified a few limitations to the software. Although the TIGER pipeline is designed to quantify 5’ and 3’ variants, it does not currently identify internal modifications, ADAR editing events, or single nucleotide polymorphisms. This feature would aid in the study of tDRs, which are heavily modified, and would potentially improve analysis of non-host sRNAs where reference genomes may be lacking. The ability to quantify internal variance is a key feature of Chimira, as well as other software, including UEA workbench^39^, and MAGI^40^. Furthermore, the TIGER pipeline does not include the analysis of PIWI-Interacting RNAs (piRNA) and a few other sRNAs, including promoter-associated sRNAs, which present unique challenges in alignments, quantification, and nomenclature^41^. Future versions of the pipeline will include less studied sRNA classes and the ability to discover new host sRNAs, as the current pipeline does not have the feature to identify novel miRNAs based on adjacent genomic sequences which is an output of other pipelines^42, 43^. Despite these limitations, the TIGER pipeline sets forth many improvements to sRNA-seq analysis.

In summary, the value of any sequencing data analysis pipeline, ultimately, is the ability to extract the most useable information from the generated data. Therefore, the goal of TIGER was to assess both host and non-host sRNAs, which greatly improved the ability to account for more reads in our sRNA-seq datasets, particularly exRNAs. TIGER also advances the field in its ability to analyze host sRNAs at the parent and fragment levels and non-host sRNAs at the genome and fragment levels. This approach may be critical to discovering novel biomarkers and intercellular communication signals that would be masked by analyzing the sRNAs by their parent RNAs. Likewise, TIGER analyzes sRNAs by class and species (genome) as well as class-independent approaches. This is very important for the study of exRNAs where the contributing parent RNA may not be annotated for the host genome, or the contributing (exogenous) species for highly abundant sRNAs may not be curated in microbiome databases. The TIGER pipeline is particularly suited for lipoprotein sRNAs which are predominantly rRNA-derived fragments of bacterial origin. Using TIGER, we were able to make critical observations comparing lipoprotein sRNAs to liver and biofluids that would not be observed by existing pipelines. Therefore, this tool is well-suited for the analysis of exRNA.

## Materials and Methods

### Animal Studies

Plasma, basal bile, urine, and livers were collected from wild-type (WT) and SR-BI-deficient (*B6;129S2-Scarb1tm1Kri/J,* SR-BI KO) mice, as previously described^44^. Mice were anesthetized with urethane (1g/kg, i.p.). The common bile duct was ligated and the gall bladder cannulated to divert bile into collection tubes. Basal bile was collected for a period of 30 min. Mice were then exsanguinated, blood was collected from the abdominal aorta in EDTA coated tubes and placed on wet ice, and tissues were dissected and snap frozen in liquid nitrogen. Plasma and tissues were stored at −80°C prior to analysis. All animal procedures were completed under active and approved IACUC protocols.

### Lipoprotein isolation

To separate HDL and apolipoprotein B (APOB)-containing lipoproteins from mouse plasma, 200 µL of 0.22-µm filtered-plasma samples were diluted to 500 µL in size-exclusion chromatography (SEC) running buffer (10 mM Tris-HCl, 0.15 M NaCl, 0.2% NaN_3_) and injected an ÄKTA SEC system (GE Healthcare) with three in-series Superdex-200 Increase gel filtration columns (10/300 GL; GE Healthcare). Samples were applied to the column with a flow rate of 0.3 mL/min at room temperature and eluate collected as 72 × 1.5 mL fractions using a F9-C 96-well plate fraction collector (GE Healthcare). Each fraction was analyzed for total protein (BCA; Pierce), total cholesterol (Raichem), and triglycerides (Raichem) to identify fractions corresponding with HDL and APOB particles. Due to the SEC set-up, we were not able to separate VLDL from LDL particles, and thus, we collected fractions covering both lipoprotein classes, referred to here as APOB. Fractions corresponding with each lipoprotein group were pooled, concentrated with Amicon Ultra-4 10 kDa centrifugal filters (Millipore) to <200 µL volume, and protein concentrations were quantified by BCA assays (Pierce). Based on the distribution of total cholesterol, triglycerides, and protein, fractions corresponding to HDL and APOB were collected, pooled, and concentrated.

### RNA Isolation

To differentiate lipoprotein sRNA signatures from liver and biofluids, and determine the impact of SR-BI-deficiency, samples were collected from *Scarb1*^*-/-*^ (SR-BI KO) and wild-type (WT) mice. Total RNA was extracted from HDL (WT N=7, SR-BI KO N=7) and APOB (WT N=7, SR-BI KO N=7) particles, as well as livers (WT N=7, SR-BI KO N=7), bile (WT N=7, SR-BI KO N=6), and urine (WT N=5, SR-BI KO N=6). RNA was isolated from equal inputs of either bile (volume), liver (mg), HDL (protein) or APOB (protein) using miRNAEasy Mini kits (Qiagen). Specifically, 30 µL of primary bile, 120 μg of APOB, 180 μg of HDL or 20 mg of liver were added to 1 mL of Qiazol. Livers were homogenized in Qiazol with High-Impact Zirconium beads using a Bead Bug Homogenizer (Benchmark Scientific). After removal of beads, subsequent steps for liver RNA extraction were followed according to manufacturer’s protocol. Bile, APOB and HDL RNA isolations were processed according to manufacturer’s protocol, except that after addition of ethanol, samples were incubated at −80°C overnight before application to isolation columns, and were eluted with a volume of 50 μL. Liver RNA samples were quantified by Take3 plates (BioTek).

### Real-Time PCR

Total RNA from equimolar amounts of HDL or APOB protein and equivolume amounts of bile or urine samples were diluted 1:10; 50 ng of total RNA from liver was used for reverse transcription using either miRCURY LNA universal RT kit (Exiqon) or TaqMan miRNA Reverse Transcription kit, as per manufacturer’s instructions. Real-time PCR was performed with the QuantStudio 12K Flex Real-Time PCR System (Life Technologies) using either: A) miRCURY LNA SYBR Green PCR kit (Exiqon) and either miRNA-specific or custom-sequence specific LNA probes (Exiqon) or B) TaqMan miRNA-specific probes. Relative quantitative values (RQV) were determined for both HDL and cellular miRNA analyses. RQV = 2^-dCt^. exo_rDR_Pflo23S 5’-AGAGAACTCGGGTGAAGGAACT-3’, exo_rDR_Vsp 5’-TGGGTGTGACGGGGAAGCAGG-3’, exo_rDR_Jliv 5’-GACCAGGACGTTGATAGGCTGGGTGTGGAAGTG-3’, miscRNA_Rpph1 5’-CGGGCCTCATAACCCAATTCAGACTACTCTCCCCCGCCCTC-3’, snDR_Gm262325’-GCGGGAAACTCGACTGCATAATTTGTGGTAGTGGGGGACTGCGTTCGCGCTCTCCCCTG-3’, snDR_Gm22866 5’-ATAATTTGTGGTAGTGGGG-3’, tDR-GluCTC5’-TCCCTGGTGGTCTAGTGGTTAGGATTCGGCGCTCTC-3’, and tDR-GlyGCC 5’-GCATTGGTGGTTCAGTGGTAGAATTCTCGC-3’. For HDL, APOB, bile, and urine samples, an arbitrary housekeeping Ct = 32 was applied, and RQVs for liver sRNAs were normalized by U6.

### Small RNA sequencing

NEXTflex Small RNA Library Preparation Kits v3 for Illumina^®^ Platforms (BioO Scientific) were used to generate cDNA libraries for sRNA-seq. Briefly, 1 µg of liver total RNA was used as input for adapter ligation, as per manufacturer’s protocol. For bile, APOB and HDL RNA, 10.5 μL (21%) of the RNA isolation eluate was used as input for adapter ligation. Library generation was performed according to manufacturer’s protocol (BioO Scientific) with a modification to the amplification step, as liver libraries received 18 cycles and bile, APOB and HDL libraries received 27 cycles. After amplification, samples were size-selected using a Pippin-Prep (Sage Science) -- set for a range of 135-200 nts in length -- and subsequently purified and concentrated using DNA Clean and Concentrator 5 kit (Zymo). Individual libraries were then screened for quality by High-Sensitivity DNA chips using a 2100 Bioanalyzer (Agilent) and quantified using High-Sensitivity DNA assays with Qubit (Life Technologies). Equal concentrations of all individual libraries were pooled for multiplex sequencing runs, and concentrated using DNA Clean and Concentrator 5 kit (Zymo). For rigor in down-stream comparisons, all 66 sequencing libraries were randomized and run independently on three individual sequencing lanes. Single-end sequencing (75 cycles) of multiplexed libraries were performed on an Illumina NextSEQ 500 at the Vanderbilt Technologies for Advanced Genomics (VANTAGE) core laboratory. Each library was sequenced at an average depth of 16.28 million reads/sample.

### Data analysis

The TIGER pipeline has many unique analysis features built into seven modules for low-level and high-level analyses with data visualization packages. The first module contains pre-processing steps (green) prior to data analysis (**Figure 1**). To assess raw data quality, FastQC was performed at the raw read level to check for base quality, total read counts, and adapter identification. Cutadapt was then used to trim 3’ adapters from processed reads (-a TGGAATTCTCGGGTGCCAAGG). Although this pipeline can analyze sRNA-seq data prepared by different library generation methods, TIGER was optimized to analyze sRNA-seq data prepared by ligation of adapters containing 4 terminal degenerate bases, which reduce ligation bias (e.g. BioO Scientific NEXTflex Small RNA-seq kit v3). Cutadapt was then used to remove the first and last 4 bases from the trimmed reads and all trimmed reads <16 nts in length were removed (-m 16 -u 4 -u −4). After trimming, read length distributions were plotted and FastQC was performed on trimmed reads to validate the efficiency of adapter trimming. The processed reads were then summarized and plotted. To generate identical read files, trimmed reads in each sample were collapsed into non-redundant “identical” reads in FASTQ format and copy numbers were recorded for downstream analysis. Preprocessed reads were then analyzed by the Host Genome & Database (blue) and Class-Independent (red) modules in parallel (**Fig.1**). In the Host Genome & Database alignment module (blue), bowtie (v1.1.2) was used to map reads to a costumed database with option (-a -m 100 --best - strata -v 1) which allows 1 mismatch (MM) and 100 multi-mapped loci, and only the best matches were recorded. The costumed database was constructed by the host genome and known sequences of host mature transcripts curated in specific library databases – tRNAs (http://gtrnadb.ucsc.edu/GtRNAdb2/) and rRNA (http://archive.broadinstitute.org/cancer/cga/rnaseqc_download). A small number of parent tRNA genes contain introns and the mature transcript differs from the genomic sequence; therefore, the incorporation of mature tRNA transcripts from GtRNAdb database into the genomic alignment overcame these limitations. This approach allows for the detection of tDRs spanning exon junctions and allows reads the chance to be mapped to other non-tRNA loci in the genome with best alignment score which reduces the false positive tDR reads that can result from database only alignment strategies. Counting and differential expression analysis of miRNAs, tDRs, rDRs, snDRs, snoDRs, and other miscellaneous sRNAs (miscRNA), including yDRs and lincDRs, were performed. The pipeline does not quantify Piwi-interacting RNA (piRNA) or circular RNAs (circRNA), but this function can be amended. All prepossessed quality reads were assigned to different classes of annotated sRNAs using distinct rules -- miRNA: 1 MM, ≥16nt, offset −2, −1, 0, 1, 2 and tDR, snDRs, snoDRs, yDRs, and miscRNAs: 1 MM, ≥16nt, overlap ≥0.9 overlap. Based on the extensive genomic coverage of lncRNAs and repetitive elements and conservation of rRNAs, the TIGER pipeline applies more stringent assignment rules for lncDRs and rDRs – perfect match, ≥20 nt, and ≥90% overlap with parent lncRNAs or rRNAs. Furthermore, reads assigned to lncDRs must only be aligned to lncRNA coordinates and not to any other loci in the genome. All reads ≥20 nts in length and not aligned to the costumed database were extracted and tested for alignment as non-host reads. After tabulation of read counts, high-end analyses were performed on host sRNAs. These include categorical analysis and visualization, principal component analysis, hierarchical clustering and correlation of samples and groups at the parent and individual fragment levels. Differential expression detection of tabulated read counts were performed by DEseq2^45^. In addition, miRNAs were analyzed at the canonical, isomiR, NTA, NTA base, and isomiR NTA levels. Non-host reads were then analyzed using the Non-Host Genome (Purple) and Non-Host Library (Gold) modules in parallel (**Fig.1**). In the Non-Host Genome module, reads were aligned in parallel to two collections of bacterial genomes: a human microbiome (HMB) collection and a hand-curated list of environmental bacteria observed during sequencing of human and mouse lipoproteins. The HMB list was compiled by reducing 3,055 bacterial genomes available from the Human Microbiome Project (www.hmpdacc.org) to single non-redundant genera, and extracting the largest available complete genome for each genera. Conversely, to generate the environmental bacteria list, the top 100 most abundant sequences in a control HDL cohort, that were not mapped to the host genome, were submitted to NCBI for BLASTn. All hits that showed 100% coverage and 100% identity were then compiled; non-redundant genera were extracted; redundant genera to the HMB were removed. Representative genomes from the remaining species were then compiled to the environmental bacteria list (ENV). Additionally, a small group of fungal genomes associated with the human pathology were also collected. The HMB, ENV, and fungal modules contain 206, 167, 8 representative genomes, respectively. Due to high conservation between bacterial genomes and multi-mapping issues, a different bowtie option (-a -m 1000 --best -strata -v 0) was used which allowed perfect match only and 1000 multi-mapped loci. Reads were aligned to the HMB, ENV, and fungal groups in parallel and, thus, the same reads could have been counted in multiple groups. The fraction of reads that align to both databases (HMB, ENV) and the reads that are unique to specific databases were plotted. Differential expression and high-end analyses, as described above, were performed at the genome level (total normalized read count for each genome) and at the individual read level. In parallel, non-host reads were also analyzed by the Non-Host Library (Gold) module where they were aligned to non-coding RNA databases with same bowtie option as non-host genome analysis. To identify possible non-host miRNAs (xenomiRs) in sRNA-seq datasets, all non-host reads were aligned perfectly to annotated miRNAs in miRBase (miRBase.org) and tabulated. Similarly, non-host reads were aligned to all tRNAs in the GtRNAdb database (GtRNAdb2). Extensive categorical analysis of parent non-host tRNAs were performed at the kingdom, genome (species), amino acid, anti-codon, and fragment (read) levels. All assigned non-host tDRs underwent differential expression analysis, high-end analysis, and data visualization, as described above. Non-host reads were also aligned to prokaryotic and eukaryotic rRNA transcripts in SILVA database (https://www.arb-silva.de). TIGER limits the analysis of non-host rDR to the kingdom level for counting, differential expression analysis and high-end analysis.

The TIGER pipeline also analyzed the top most abundant reads independent of class or annotation in parallel of the host genome, non-host genome, and database modules. The Class-Independent module (red) ranked and filtered the top 100 most abundant reads in each sample independent of genomic annotation. The list of top 100 reads from all samples were combined, a count file table was generated and top 100 overall reads were used to perform hierarchical clustering and correlations at the individual sample and group levels. Differential expression analyses were performed by DEseq2, and significantly altered sequences were searched in NCBI nucleotide database using BLASTn to identify possible sources (species). All results from the host genome, class-independent, non-host genome, and non-host database modules were then analyzed by the Summary & Data Visualization (dark blue) module (**Fig.1**). In this module, TIGER summarized and organized many of the individual comparisons. For example, individual volcano plots were graphed into larger matrices grouping different classes of sRNAs and/or genomic groups (e.g. bacteria and fungi). This module also generated a comprehensive table for all mapped reads listing the assignments for each read across modules. Moreover, positional coverage of sRNAs against host parent RNAs were plotted for miRNAs, tDRs, snDRs, and rDRs. Positional base coverages were also plotted for individual samples, groups, and significantly altered tDRs and snDRs. For groups, the means of normalized positional coverage counts (base positional counts per million mapped total reads) for individual samples in the groups were plotted. Furthermore, this module identified sRNA classes and genomes for the top 100 ranked reads (analyzed earlier in the Class-Independent module) and graphed the linkages by circos plots. Finally, this module summarized the read counts in each task and determined the fraction of total reads that were assigned to any module, genome, or database. For example, pie charts and stacked bar charts were generated to illustrate the fraction of reads mapped to the host genome and non-host genome and the fraction of unmapped reads. All unmapped and unaccounted for reads entered the Final Unmapped Analysis (orange) module (**Fig.1**). In this module, the top 100 analysis was reapplied to all unmapped and unaccounted reads, as described above. After ranking, filtering, and tabulation, differential expression analysis was performed and the significantly altered unmapped reads were searched in BLASTn to identify possible genomes not contained in the TIGER analysis. These unique features were designed to extensively and exhaustively analyze sRNA-seq data on lipoproteins (e.g. HDL and apoB particles) and extracellular fluids (e.g. bile and urine) which have many different types of sRNAs and diverse species.

### Data Visualization

Read counts were reported as both raw counts and normalized count per million total counts (RPM). RPMs were used for stacked bar plots in each module. Cluster analysis were performed and visualized by heatmap3^46^. Principle component analysis were performed based on normalized expression value calculated by the variance stabilizing transformation in DESeq2. DESeq2 was used to perform miRNA, tDR and other sRNA differential expression analyses. Significantly differential expressed sRNAs with adjusted p-value less <0.05 and absolute fold change >1.5 will be highlighted in volcano plot (red, increased; blue, decreased) and outputted as tabulated file for further validation. Differential expression results were plotted as volcano plot, venn diagram, and heatmap. Categorical analyses of tDRs based on amino acid and anti-codons of the parent tRNAs were also quantified and plotted. Likewise, categorical analysis of snDRs based on U class were analyzed and plotted. Non-metric multidimensional scaling of Bray-Curtis dissimilarity indexes, homogeneity analysis of group dispersions, and principal coordinate analysis visualization was performed using R package “vegan”. R Packages ggplot2, vegan, ggraph, igraph, reshape2, data.table, RColorBrewer, circlize, ggtern, and XML were used for data visualization.

### Statistics

For continuous variables, mean and standard error of the mean (S.E.M.) were used. Comparisons with two variables were calculated using Welch two sample t-tests, two-way Student’s t-tests, or Mann-Whitney nonparametric tests. For comparisons with more than two variables, linear one-way analysis of variance (ANOVA) were used. Spearman ranked method was used for calculating the correlation coefficient (R). Two-sided p value <0.05 was considered statistically significant. Statistical analyses were performed using R version 3.4.3.

## Acknowledgments

We would like to acknowledge Carolin Besenboeck for her assistance in RNA methods.

## References

1. Wang K, Li H, Yuan Y, Etheridge A, Zhou Y, Huang D, Wilmes P, and Galas D., The complex exogenous RNA spectra in human plasma: an interface with human gut biota? PLoS One. 2012;7:e51009.

2. Beatty M, Guduric-Fuchs J, Brown E, Bridgett S, Chakravarthy U, Hogg RE, and Simpson DA., Small RNAsfrom plants, bacteria and fungi within the order Hypocreales are ubiquitous in human plasma. BMC Genomics. 2014;15:933.

3. Quintana JF, Makepeace BL, Babayan SA, Ivens A, Pfarr KM, Blaxter M, Debrah A, Wanji S, Ngangyung HF, Bah GS, Tanya VN, Taylor DW, Hoerauf A, and Buck AH., Extracellular Onchocerca-derived small RNAs in host nodules and blood. Parasit Vectors. 2015;8:58.

4. Vickers KC, Sethupathy P, Baran-Gale J, and Remaley AT. Complexity of microRNA function and the role of isomiRs in lipid homeostasis. J Lipid Res. 2013;54:1182–91.

5. Scott DD and Norbury CJ. RNA decay via 3’ uridylation. Biochim Biophys Acta. 2013;1829:654–65.

6. Knouf EC, Wyman SK, and Tewari M., The human TUT1 nucleotidyl transferase as a global regulator of microRNA abundance. PLoS One. 2013;8:e69630.

7. Bartel DP. MicroRNAs: genomics, biogenesis, mechanism, and function. Cell. 2004;116:281–97.

8. Vitsios DM and Enright AJ. Chimira: analysis of small RNA sequencing data and microRNA modifications. Bioinformatics. 2015;31:3365–7.

9. Vickers KC, Roteta LA, Hucheson-Dilks H, Han L, and Guo Y., Mining diverse small RNA species in the deep transcriptome. Trends in biochemical sciences. 2015;40:4–7.

10. Chen CJ and Heard E.Small RNAs derived from structural non-coding RNAs. Methods. 2013;63:76–84.

11. Li Z, Ender C, Meister G, Moore PS, Chang Y, and John B., Extensive terminal and asymmetric processing of small RNAs from rRNAs, snoRNAs, snRNAs, and tRNAs. Nucleic Acids Res. 2012;40:6787–99.

12. Vickers KC, Roteta LA, Hucheson-Dilks H, Han L, and Guo Y., Mining diverse small RNA species in the deep transcriptome. Trends in biochemical sciences. 2015;40:4–7.

13. Baras AS, Mitchell CJ, Myers JR, Gupta S, Weng LC, Ashton JM, Cornish TC, Pandey A, and Halushka MK. miRge - A Multiplexed Method of Processing Small RNA-Seq Data to Determine MicroRNA Entropy. PLoS One. 2015;10:e0143066.

14. Capece V, Garcia Vizcaino JC, Vidal R, Rahman RU, Pena Centeno T, Shomroni O, Suberviola I, Fischer A, and Bonn S.Oasis: online analysis of small RNA deep sequencing data. Bioinformatics. 2015;31:2205–7.

15. Boon RA and Vickers KC. Intercellular transport of microRNAs. Arterioscler Thromb Vasc Biol. 2013;33:186–92.

16. Vickers KC and Remaley AT. Lipid-based carriers of microRNAs and intercellular communication. Curr Opin Lipidol. 2012;23:91–7.

17. Vickers KC, Palmisano BT, Shoucri BM, Shamburek RD, and Remaley AT. MicroRNAs are transported in plasma and delivered to recipient cells by high-density lipoproteins. Nat Cell Biol. 2011;13:423–33.

18. Cloonan N, Wani S, Xu Q, Gu J, Lea K, Heater S, Barbacioru C, Steptoe AL, Martin HC, Nourbakhsh E, Krishnan K, Gardiner B, Wang X, Nones K, Steen JA, Matigian NA, Wood DL, Kassahn KS, Waddell N, Shepherd J, Lee C, Ichikawa J, McKernan K, Bramlett K, Kuersten S, and Grimmond SM. MicroRNAs and their isomiRs function cooperatively to target common biological pathways. Genome Biol. 2011;12:R126.

19. Neilsen CT, Goodall GJ, and Bracken CP. IsomiRs–the overlooked repertoire in the dynamic microRNAome. Trends Genet. 2012;28:544–9.

20. Vickers KC, Sethupathy P, Baran-Gale J, and Remaley AT., The Complexity of microRNA Function and the Role of IsomiRs in Lipid Homeostasis. J Lipid Res. 2013.

21. Baran-Gale J, Fannin EE, Kurtz CL, and Sethupathy P.Beta cell 5’-shifted isomiRs are candidate regulatory hubs in type 2 diabetes. PLoS One. 2013;8:e73240.

22. Burroughs AM, Ando Y, de Hoon MJ, Tomaru Y, Nishibu T, Ukekawa R, Funakoshi T, Kurokawa T, Suzuki H, Hayashizaki Y, and Daub CO. A comprehensive survey of 3’ animal miRNA modification events and a possible role for 3’ adenylation in modulating miRNA targeting effectiveness. Genome Res. 2010;20:1398–410.

23. Koppers-Lalic D, Hackenberg M, Bijnsdorp IV, van Eijndhoven MAJ, Sadek P, Sie D, Zini N, Middeldorp JM, Ylstra B, de Menezes RX, Wurdinger T, Meijer GA, and Pegtel DM., Nontemplated nucleotide additions distinguish the small RNA composition in cells from exosomes. Cell reports. 2014;8:1649–1658.

24. Selitsky SR and Sethupathy P.tDRmapper: challenges and solutions to mapping, naming, and quantifying tRNA-derived RNAs from human small RNA-sequencing data. BMC Bioinformatics. 2015;16:354.

25. Kaczor-Urbanowicz KE, Kim Y, Li F, Galeev T, Kitchen RR, Koyano K, Jeong SH, Wang X, Elashoff D, Kang SY, Kim SM, Kim K, Kim S, Chia D, Xiao X, Rozowsky J, and Wong DTW., Novel approaches for bioinformatic analysis of salivary RNA Sequencing data in the biomarker development process. Bioinformatics. 2017.

26. Acton S, Rigotti A, Landschulz KT, Xu S, Hobbs HH, and Krieger M., Identification of scavenger receptor SR-BI as a high density lipoprotein receptor. Science. 1996;271:518–20.

27. Zhang Y, Silva JR Da, Reilly M, Billheimer JT, Rothblat GH, and Rader DJ., Hepatic expression of scavenger receptor class B type I (SR-BI) is a positive regulator of macrophage reverse cholesterol transport in vivo. J Clin Invest. 2005;115:2870–4.

28. Wiersma H, Gatti A, Nijstad N, Oude Elferink RP, Kuipers F and Tietge UJ. Scavenger receptor class B type I mediates biliary cholesterol secretion independent of ATP-binding cassette transporter g5/g8 in mice. Hepatology. 2009;50:1263–72.

29. Zanoni P, Khetarpal SA, Larach DB, Hancock-Cerutti WF, Millar JS, Cuchel M, DerOhannessian S, Kontush A, Surendran P, Saleheen D, Trompet S, Jukema JW, De Craen A, Deloukas P, Sattar N, Ford I, Packard C, Majumder A, Alam DS, Di Angelantonio E, Abecasis G, Chowdhury R, Erdmann J, Nordestgaard BG, Nielsen SF, Tybjaerg-Hansen A, Schmidt RF, Kuulasmaa K, Liu DJ, Perola M, Blankenberg S, Salomaa V, Mannisto S, Amouyel P, Arveiler D, Ferrieres J, Muller-Nurasyid M, Ferrario M, Kee F, Willer CJ, Samani N, Schunkert H, Butterworth AS, Howson JM, Peloso GM, Stitziel NO, Danesh J, Kathiresan S, Rader DJ, Consortium CHDE, Consortium CAE, and Global Lipids Genetics C. Rare variant in scavenger receptor BI raises HDL cholesterol and increases risk of coronary heart disease. Science. 2016;351:1166–71.

30. Varban ML, Rinninger F, Wang N, Fairchild-Huntress V, Dunmore JH, Fang Q, Gosselin ML, Dixon KL, Deeds JD, Acton SL, Tall AR, and Huszar D., Targeted mutation reveals a central role for SR-BI in hepatic selective uptake of high density lipoprotein cholesterol. Proc Natl Acad Sci U S A. 1998;95:4619–24.

31. Zhong CY, Sun WW, Ma Y, Zhu H, Yang P, Wei H, Zeng BH, Zhang Q, Liu Y, Li WX, Chen Y, Yu L, and Song ZY. Microbiota prevents cholesterol loss from the body by regulating host gene expression in mice. Scientific reports. 2015;5:10512.

32. Zhang X, Cozen AE, Liu Y, Chen Q, and Lowe TM., Small RNAModifications: Integral to Function and Disease. Trends in molecular medicine. 2016;22:1025–1034.

33. Telonis AG, Loher P, Kirino Y, and Rigoutsos I.Consequential considerations when mapping tRNA fragments. BMC Bioinformatics. 2016;17:123.

34. Wei Z, Batagov AO, Schinelli S, Wang J, Wang Y, Fatimy R El, Rabinovsky R, Balaj L, Chen CC, Hochberg F, Carter B, Breakefield XO, and Krichevsky AM., Coding and noncoding landscape of extracellular RNA released by human glioma stem cells. Nature communications. 2017;8:1145.

35. Yeri A, Courtright A, Reiman R, Carlson E, Beecroft T, Janss A, Siniard A, Richholt R, Balak C, Rozowsky J, Kitchen R, Hutchins E, Winarta J, McCoy R, Anastasi M, Kim S, Huentelman M, and Van Keuren-Jensen K.Total Extracellular Small RNA Profiles from Plasma, Saliva, and Urine of Healthy Subjects. Scientific reports. 2017;7:44061.

36. Quinn JF, Patel T, Wong D, Das S, Freedman JE, Laurent LC, Carter BS, Hochberg F, Van Keuren-Jensen K, Huentelman M, Spetzler R, Kalani MY, Arango J, Adelson PD, Weiner HL, Gandhi R, Goilav B, Putterman C, and Saugstad JA., Extracellular RNAs: development as biomarkers of human disease. Journal of extracellular vesicles. 2015;4:27495.

37. Zernecke A and Preissner KT. Extracellular Ribonucleic Acids (RNA) Enter the Stage in Cardiovascular Disease. Circ Res. 2016;118:469–79.

38. Willeit P, Skroblin P, Moschen AR, Yin X, Kaudewitz D, Zampetaki A, Barwari T, Whitehead M, Ramirez CM, Goedeke L, Rotllan N, Bonora E, Hughes AD, Santer P, Fernandez-Hernando C, Tilg H, Willeit J, Kiechl S, and Mayr M., Circulating MicroRNA-122 Is Associated With the Risk of New-Onset Metabolic Syndrome and Type 2 Diabetes. Diabetes. 2017;66:347–357.

39. Stocks MB, Moxon S, Mapleson D, Woolfenden HC, Mohorianu I, Folkes L, Schwach F, Dalmay T, and Moulton V., The UEA sRNA workbench: a suite of tools for analysing and visualizing next generation sequencing microRNA and small RNA datasets. Bioinformatics. 2012;28:2059–61.

40. Kim J, Levy E, Ferbrache A, Stepanowsky P, Farcas C, Wang S, Brunner S, Bath T, Wu Y and Ohno-Machado L.MAGI: a Node.js web service for fast microRNA-Seq analysis in a GPU infrastructure. Bioinformatics. 2014;30:2826–7.

41. Agirre E and Eyras E.Databases and resources for human small non-coding RNAs. Hum Genomics. 2011;5:192–9.

42. An J, Lai J, Lehman ML, and Nelson CC. miRDeep*: an integrated application tool for miRNA identification from RNA sequencing data. Nucleic Acids Res. 2013;41:727–37.

43. Friedlander MR, Chen W, Adamidi C, Maaskola J, Einspanier R, Knespel S, and Rajewsky N.Discovering microRNAs from deep sequencing data using miRDeep. Nat Biotechnol. 2008;26:407–15.

44. Wang Y, Liu X, Pijut SS, Li J, Horn J, Bradford EM, Leggas M, Barrett TA, and Graf GA., The combination of ezetimibe and ursodiol promotes fecal sterol excretion and reveals a G5G8-independent pathway for cholesterol elimination. J Lipid Res. 2015;56:810–20.

45. Love MI, Huber W, and Anders S., Moderated estimation of fold change and dispersion for RNA-seq data with DESeq2. Genome Biol. 2014;15:550.

46. Zhao S, Guo Y, Sheng Q, and Shyr Y. Advanced heat map and clustering analysis using heatmap3. BioMed research international. 2014;2014:986048.

